# Differential encoding of social identity, valence and unfamiliarity in the amygdala and piriform cortex

**DOI:** 10.1101/2025.07.04.663165

**Authors:** Cristina Mazuski, Lars-Lennart Oettl, Chenyue Ren, John O’Keefe

## Abstract

To appropriately respond to social situations, animals must rapidly identify conspecifics as individuals, gauge their familiarity and recall any good or bad associations. In previous work (Mazuski & O’Keefe, 2022), we characterized populations of neurons in the basolateral amygdala complex (BLA) that were strongly tuned to social conspecifics. In this current study, we asked whether these or other populations in the BLA encode properties necessary for social interaction: individual recognition, familiarity detection and reward association. In addition to cells in the rat lateral amygdala (LA) and basolateral amygdala (BLA), we recorded from cells in the anatomically-related piriform cortex (PIR) to see how much of the amygdala response was inherited from this input. Here, we show that these three interrelated areas encode distinct features of social conspecifics at the population and single neuron level.

We recorded from large populations of neurons using 4-shank Neuropixels probes while rats learned a social discrimination task, where one male conspecific was paired with a food reward (S+) and the other its absence (S-). For comparison, two unfamiliar animals were presented on occasional probe trials. After successful discrimination learning, we looked for transfer effects to our previously studied task where implanted rats freely interacted with all four conspecifics in an open-field environment. The LA dynamically encoded social reward learning. Initially, LA neuronal population activity could not distinguish between the S+ and S-conspecifics, but after learning distinct representations emerged. In contrast, the PIR showed no effect of social reward learning. Single PIR units encoded social identity with strong tuning to individual conspecifics both before and after learning. The BLA encoded both reward learning and social identity suggesting this is where the PIR and LA information streams converge. Unexpectedly, the BLA and LA strongly coded for the unfamiliarity of the rat – single neurons responded to the probe trial animals with firing rates 2-5x higher than to the task animals. This cross-region encoding of identity, reward and unfamiliarity was reactivated during social interaction in the new open-field environment, demonstrating that the learning was not context specific. The results throw light on how distinct neural circuits contribute to social recognition and memory and how they interact with each other.

## Introduction

Social interaction is a natural behavior with both innate and learned dimensions^1^. Social species like rats can quickly identify conspecifics using sensory cues ^2^ such as olfaction ^3^, but must flexibly adapt their behaviors to individual conspecifics on the basis of past experiences^23^. How neurons downstream of the primary sensory circuits flexibly encode social conspecifics is poorly understood.

The amygdala and piriform cortex (PIR) are two structures located in the limbic area, historically named for its location on the rim of the cortical mantle and subsequently implicated in memory, motivation and emotion^4^. Receiving direct projections from the olfactory bulb, neuronal ensembles in the PIR primarily encode odor identity^5^, but have recently been shown to encode other contextual information like spatial location^6^ and learning^7^. Located medially adjacent to the posterior PIR, the lateral (LA) and basolateral amygdala (BLA) receive multi-sensory input^8^ from multiple sensory cortices including the PIR^9^. Neurons in the LA and BLA respond to a wide range of simple and complex stimuli including but not limited to auditory tones^10,11^, different foods^12,13^, noxious/painful stimuli^14^, and social conspecifics^13,15^. It has been proposed that the LA and BLA encode valence^16^, state^17^, and concepts^18^, but a unifying theory explaining the diversity of responses observed in the amygdala has so far been elusive.

Understanding the unique contribution of the LA, BLA and PIR in social behavior is challenging because of their high levels of interconnectedness. Manipulation of these areas through lesions, chemogenetic and optogenetic activation and inhibition can affect sociability^19^, social memory^20^, social novelty^21^, and observational learning^22^, indicating that the amygdala and piriform are part of circuitry involved in social learning and memory.

However, it is difficult to interpret to what extent these results are driven by neurons of the LA, BLA and PIR or through changes in the broader social circuitry, which includes downstream areas like the prefrontal cortex and hippocampus. To address this problem, we recorded single-unit activity in the LA, BLA and PIR during social learning and memory.

In previous work^13^, we showed that BLA neurons represented salient stimuli including conspecifics during naturalistic exploration. We characterized two populations of neurons – event-specific pyramidal neurons that were highly tuned to a single class of conspecifics (e.g. male or female) and a population of broadly tuned neurons that responded to conspecifics and other salient stimuli like food. In this previous study, we did not detect any changes in firing due to the identity of an individual (e.g. one specific male versus another), and we did not systematically test how experience changed this social representation. In this current study, we addressed these questions by measuring large-scale neuronal activity in the LA, BLA and PIR during social reward learning and its subsequent reactivation during naturalistic social interaction.

To this aim, we devised a go/no-go operant task inspired by two lines of research – social recognition^23^ and associative learning^24^. Across a single session, test rats performed a task where they were presented with one of 4 rats. Two male conspecifics were presented frequently with one associated with reward (S+ stimulus rat) and the other not (S-stimulus rat). The third and fourth conspecifics (one male and one female) were presented infrequently and neither associated with reward. After social reward learning, the trained rats were allowed to engage in naturalistic unrestrained social interaction with each of the same 4 conspecifics. This experimental design allowed us to ask 1) whether these brain regions can discriminate between different conspecifics, 2) how this representation changes with valence learning, and 3) how responses during this operant task correlate with naturalistic social interactions. Our results suggest that the LA, BLA and PIR differentially encode social identity, valence and unfamiliarity, each carrying out a unique role in social cognition.

## Results

### Rats learn to associate conspecifics with reward in a social reward task

Before electrophysiological recording, male test rats were trained on a social reward go/no-go task (Fig 1A). They learned to associate one male stimulus conspecifics with a food reward (S+) or the other with the absence of all reward (S-). Test rats were trained in daily sessions of 120 trials until they reached 75% accuracy in a single session (Fig. 1A). On discrimination trials, the test rat initiated each trial by breaking an infrared sensor beam at the sampling port which opened an opaque sliding door to present one of the 2 stimulus conspecifics. The rats were physically separated by transparent walls with small holes allowing air flow but not physical contact. The test rat was required to sample the stimulus conspecific odor for a minimum of 1s before making a choice. A correct choice consisted of breaking an infrared beam at the food port within 5s of the end of the sensory sampling period for the S+ and not poking the food port when presented with the S-(Fig 1B, FigS1A-B and Videos1-2).

**Figure 1.**
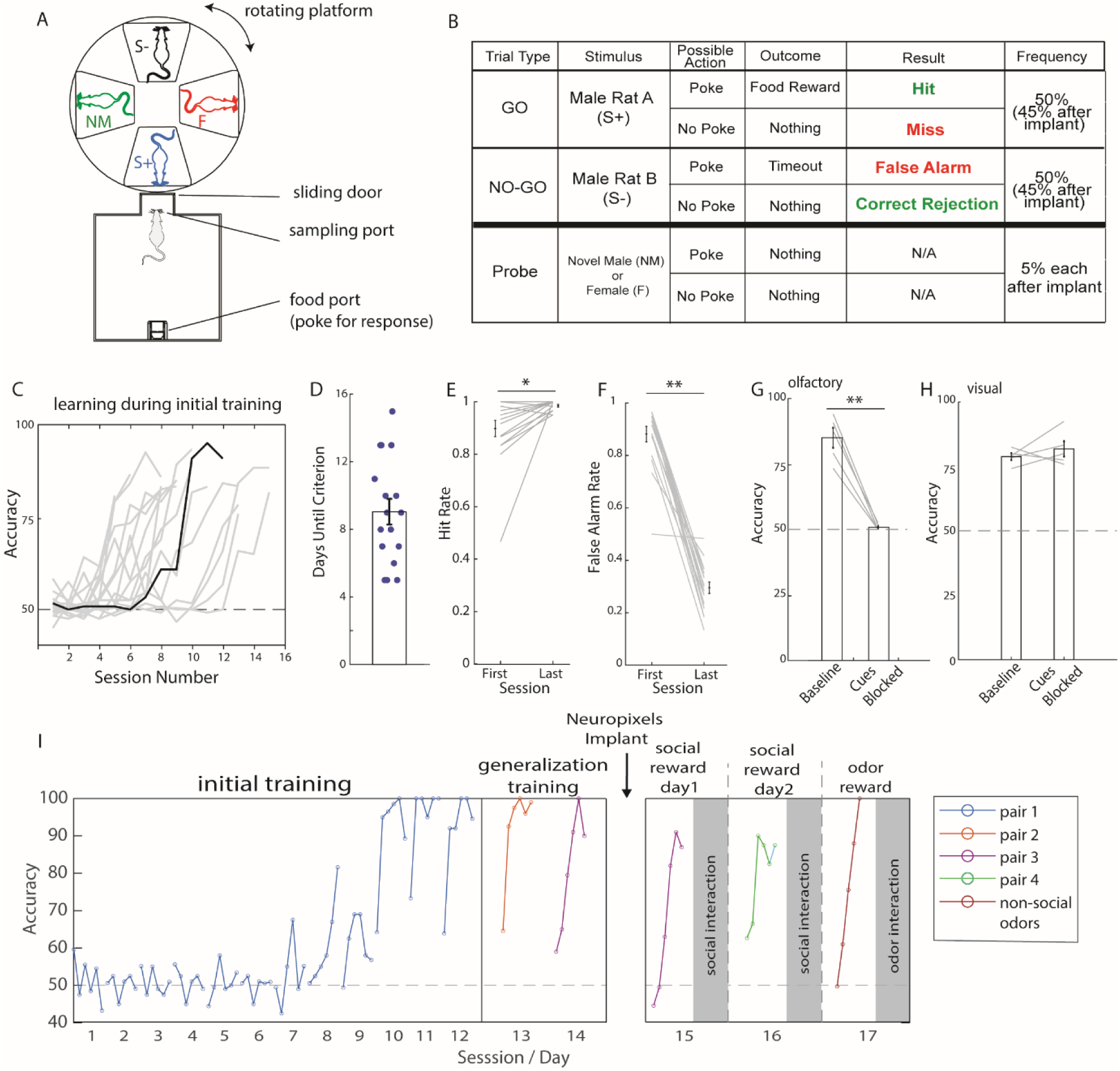
Rats can learn conspecific-valence association. (A) Diagram depicting the social olfactory go/no-go setup. (B) The full breakdown of all task possibilities. To correctly perform the task, rats must poke the food port when presented with the S+ and inhibit this behavior with the S-. (C) Individual rats’ performance (light gray) with one representative rat highlighted in black. (D) On average, rats reached criterion in 9 days **±** 0.75. (N=17 rats). (E) Rats increased hit rate (0.89 **±** 0.03 vs. 0.98 **±** 0.01, first vs. last session, *, N=17) and (F) markedly decreased false alarm rate (0.88 **±** 0.02 vs. 0.29 **±** 0.02, first vs. last session, **, N=17). (G) When olfactory cues were blocked in a subset of rats, performance dropped to chance levels (85 **±** 3.8 vs. 50.83 +-0.52, baseline vs. cues blocked, **, N=5). (H) In contrast when visual cues were blocked, performance was unaffected (78.83 **±** 1.35 vs. 81.83 +-2.95, baseline vs. cues blocked, p=0.44, N=5). (I) A diagram depicting the experimental design and performance of the representative rat (also highlighted in 1C) before and after electrophysiological recording. Every dot corresponds to 20 trials. Note the rapid relearning on each session even after reaching criterion (sessions 10 onward). Paired t-test for all comparisons, *p<0.05, **p<0.01. All values mean **±** SEM.

On average, rats learned the task within 9 days of training (Fig 1C-D), primarily by learning to not approach the food port after being presented with the S-, as evidenced by the large decrease in the false alarm rate compared to the modest increase in the hit rate across learning (Fig 1E-F, FigS1C-D). Test rats spent more time sampling the S-compared to the S+, an effect that was significant even from the initial training sessions and before significant valence learning (FigS1E), suggesting that they already could tell one animal from another and therefore that discrimination of social *identity* and social *valence* are dissociable in this task. Surprisingly, even after reaching criterion, the animals often started each day’s learning at a level close to 50% but rapidly attained a high-performance level in a small number of trials (Fig 1I).

As expected from the design of the task, we confirmed that rats primarily used olfactory cues to perform the discrimination. When we blocked active airflow from the stimulus to test rat, performance dropped to chance level (Fig 1G). In contrast, when visual cues were blocked by replacing the transparent walls with opaque walls that still permitted airflow (Fig 1H), rats were still able to perform the task.

After initial training, learning quickly generalized to new pairs of stimulus conspecifics (Fig 1I and S1F). After rats successfully generalized to a minimum of 2 new stimulus pairs, they were implanted with 4-shank Neuropixels probes targeting the posterior ventrolateral brain (Fig 1I, 2A and FigS2).

In total, we recorded electrophysiological data from 10 rats during the social reward task (350±69 neurons per rat, mean±SEM). During recording, the rats were also presented with infrequent probe trials of 2 conspecifics not relevant to the task (Fig 1B, 1I). Performance on standard trials was not affected by surgery or these probe trials. Rats spent increased time sampling the probe conspecifics, suggesting that they could detect the difference between them and the usual task conspecifics (FigS1G and Video3-4).

### Social identity and social valence are represented differently in the amygdala and piriform cortex

We recorded from hundreds of neurons in the lateral amygdala (LA, N=380), basolateral amygdala (BLA, N=416) and posterior piriform cortex (PIR, N=1221) (Fig 2A and Fig S2) during the social reward task. As rats showed rapid relearning during each single-day session, we calculated the learning inflection point from the behavioral performance of each session and divided the data into pre-learning and post-learning trials (Fig 2B). The results from the 2 days of social reward learning were comparable (Fig S3), and were combined for subsequent analyses.

**Figure 2.**
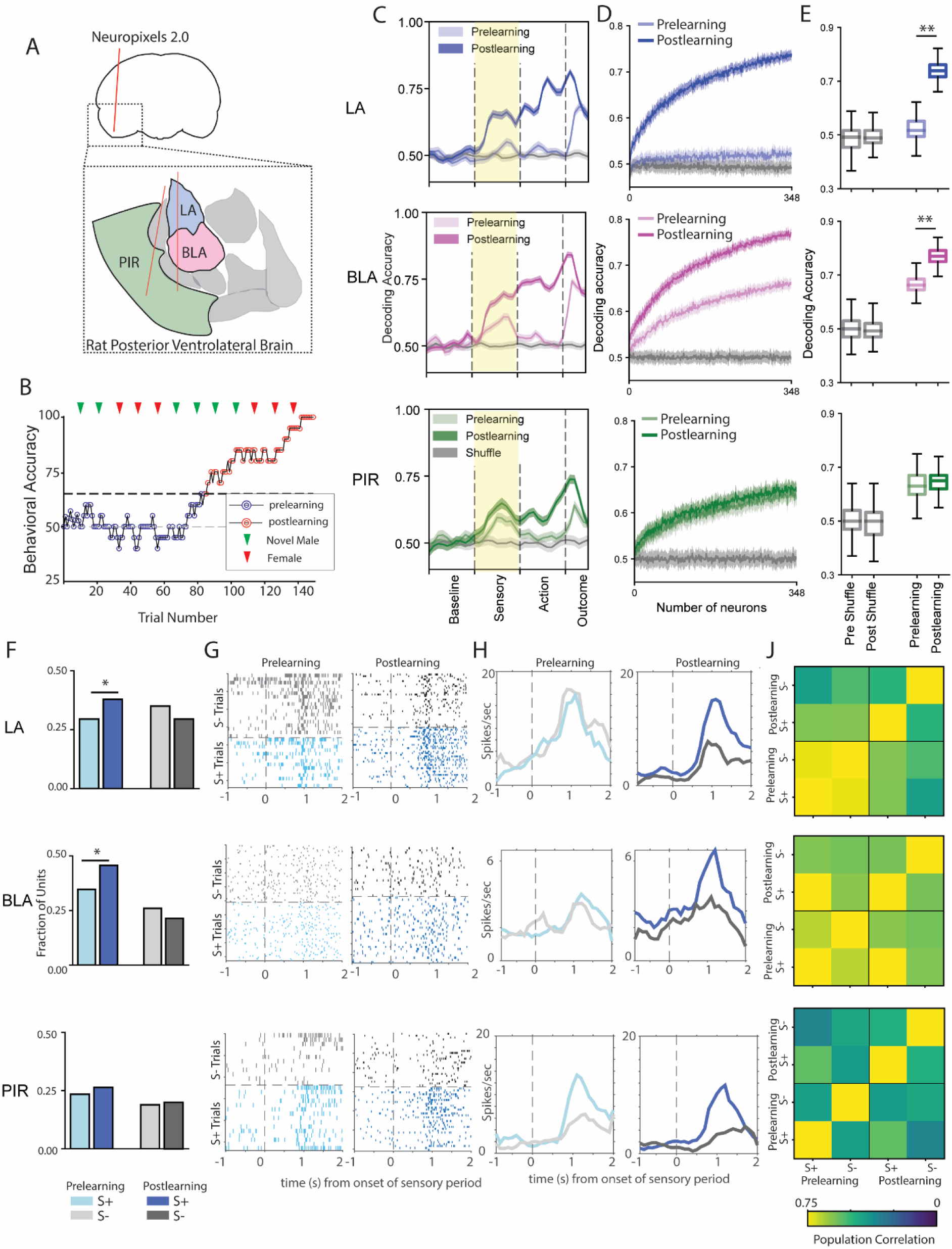
The amygdala and piriform cortex display distinct learning dynamics during social reward learning. (A) Chronically implanted Neuropixels 2.0 probes targeted to the posterior ventrolateral brain to record from the LA, BLA and PIR. Two example probe tracts shown in orange. (B) A representative example of behavioral performance during a single-day. Based on the learning inflection point (black dotted line), each session was divided into pre- and post-learning trials. Probe-trial presentations of a novel male (green carrot) or female (red carrot) conspecific occurred every 10 trials in a randomized fashion. (C) Conspecific identity (S+ vs. S-) was reliably decoded from population neural activity in all 3 regions (LA: top, BLA: middle, PIR: bottom, support vector machine (SVM) classifier trained on 50 units per region, N=100 iterations). However different dynamics emerge during the sensory sampling period (shaded yellow). The LA and BLA showed large improvements in decoding during this period, while the PIR was able to decode conspecific identity before learning and this did not improve significantly with learning. (D) We trained a SVM classifier with increasing number of neurons per region (from 2 to 348) to calculate decoding accuracy during the sensory period. (E) Quantification of decoding performance from the maximum number of neurons (348). All regions show significant decoding of task animal identity compared to shuffled data. A repeated-measures ANOVA revealed significant main effects of learning condition (i.e. shuffled, pre-, post-learning, F(3, 1188) = 17,059, p < .001) and brain region, F(2, 1188) = 1179, p < .001, as well as a significant condition x region interaction, F(6, 1188) = 1120. Post-hoc comparisons showed significant learning-related increases in decoding in the LA (*pre:* 0.52±0.004; *post:* 0.74±0.003, p<0.001) and BLA (*pre:* 0.66±0.003, *post:* 0.77±0.003, p<0.001), but not the PIR (*pre:* 0.63±0.005, *post:* 0.65±0.005, p=0.11) N=100 iterations. **F**) In LA and BLA, the fraction of units exhibiting significant responses to the S+ increased while the number of units responding to the S-modestly decreased. PIR did not change with learning (change in S+ responsive: LA: χ2(1, N=380) = 5.66, *p<0.05; BLA: χ2(1, N=416) = 10.13, *p<0.05; PIR: χ2(1, N=1221) = 2.68, p=0.10, change in S-responsive: LA: χ2(1, N=380) = 2.41, p=0.12; BLA: χ2(1, N=416) = 2.15, p=0.14; PIR: χ2(1, N=1221) = 0.32, p=0.57) **(G)** Representative single units firing (LA: top, BLA: middle, PIR: bottom) during individual trials is shown in rasterplots and **(H)** average firing rate across trial type. **(J)** Correlation of the pre- and post-learning population activity for all responsive neurons reveals social reward learning in LA (top), social identity encoding in PIR (bottom) and a mixed representation in BLA (middle).

To compare neuronal data across different trial lengths, we interpolated the spike times to align neuronal data to behavioral epochs (sensory, action and outcome epochs, see methods and S1B). We tested whether conspecific identity (S+ or S-) could be decoded from population activity before and after learning (Fig 2C). During the sensory sampling period, we found striking differences across regions in how they encode social identity versus social valence. Decoding accuracy of social valence or reward-learning induced changes was only present in the amygdala (LA and BLA, Fig. 2D-E). In contrast, neural activity in the PIR and BLA, but not the LA, reliably decoded social identity even before learning occurred.

At the level of single-units, there were valence-induced changes during the sensory period in both LA and BLA. Social reward learning resulted in a significant increase in the fraction of units responding to the S+ and a small decrease in S-units in both LA and BLA (Fig 2F). No such change was observed in PIR. In addition, single-neuron firing rate activity changed in response to reward learning only in LA and BLA, but not PIR where, in contrast, activity was tuned to individual conspecifics even before any learning took place. Activity in some PIR neurons remapped over time (see Fig 2J and S4A). We correlated population activity of responsive neurons per region during the sensory period and found that the strong correlation between the pre-learning representation of the S+ and S- in the LA and BLA decreased with learning suggesting that the population activity became more distinct. In contrast, the representation of conspecifics in the PIR was distinct before learning and remained distinct after (Fig 2J). In all regions, population activity could decode stimulus identity across contexts (pre-vs. post-learning) but this was weaker than within-context decoding accuracy (Fig S4B-C).

### Unfamiliarity is strongly encoded in the amygdala

During social reward learning, we presented either a novel male (NM) or a female (F) conspecific in probe trials that occurred every 10 trials. Single-units within the LA, BLA and PIR responded strongly to the probe trial animals, oftentimes at firing rates 2-5x higher than for the task animals (Fig 3A and S5C). We compared the neural representation of the 4 conspecifics (S+, S-, NM and F) by plotting the population neuronal trajectory during the sensory period for each (Fig S5A). While all three regions had distinct representations of the four conspecifics, the LA and BLA showed a strong divergence in how the task and probe trial animals were encoded. In contrast in the PIR, all four animals were represented with distinct but equally spaced trajectories (Fig S5B).

**Figure 3.**
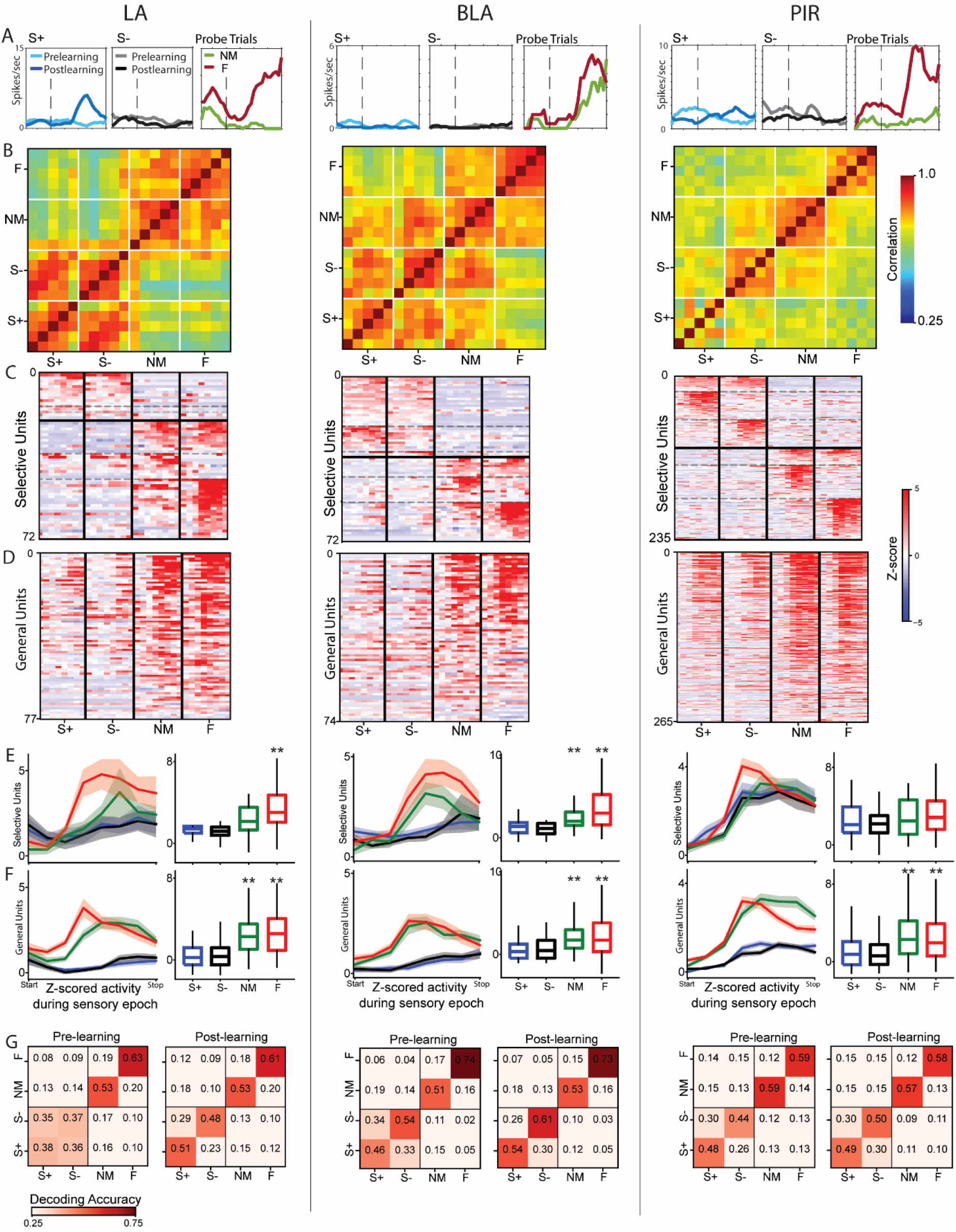
LA and BLA show strong preferences for the unfamiliar animals while PIR responds similarly to all 4 animals. **(A)** Representative single units from the LA, BLA, and PIR (left, middle, right) showed markedly stronger responses (2-5x) to probe trial animals compared to task animals. These responses were seen for both the female and novel male (FigS5C) **(B)** Population correlation of sensory responses from the first five presentations of each animal revealed distinct representation of individuals in the PIR, while population response in the LA and BLA was organized by animal class (task vs. probe) and in the BLA also by sex (M vs. F). **(C)** Z-scored neuronal responses during the first 5 trials of each animal were sorted based on their selectivity. Selective units tuned to either task or probe trial animals were sorted by preferred firing (both task animals, S+ only, S-only, both probe trial animals, NM only, F only). **(D)** Broadly tuned neurons that responded to both task and probe trial animals were sorted by average response strength. **(E)** Mean z-scored sensory responses for selective neurons (from C) showed significantly stronger responses to probe animals in the LA and BLA (**p < 0.01). The PIR showed no significant difference (p=0.29, Kruskal-Wallis test with Bonferroni-corrected post-hoc comparisons). The LA had significantly stronger responses to the female, while the BLA had stronger response to both the NM and F compared to task animals. **(F)** General neurons (from D) showed significantly stronger firing to the probe trial animals (**p<0.01) in all three regions. (G) Decoding accuracy of the 4 animals shows distinct dynamics across regions and in response to learning. The LA (left) is unable to discriminate task animals prior to learning, but shows consistently strong discrimination of non-task animals. In contrast, the PIR (right) can reliably distinguish between the 4 animals before learning takes place and this does not change with learning. The BLA (middle) exhibits a mixture of dynamics with some identity encoding similar to the piriform and substantial social reward learning.

To examine whether the differences in conspecific representation in the LA and BLA was driven by novelty or presentation number, we correlated the population sensory response from responsive units for the first 5 presentations of each animal (Fig 3B). In the amygdala regions, we see higher correlations between presentations of animals from the same class (i.e. task animals or probe trial animals) than across classes. In contrast, the PIR has distinct population representations for all four animals. These results suggest that in the amygdala, novelty alone cannot explain the difference in neuronal tuning. Instead, it is likely that the LA and BLA encode a more abstract concept like the level of social unfamiliarity or unexpectedness.

The difference in representation of the task and probe trial animals in the LA and BLA occurred at the level of single neurons. We sorted individual units by their tuning selectivity to task or probe trial animals and found that single-units within the LA and BLA showed more selectivity to probe trial animals (Fig 3C-D) and stronger responses (Fig 3E-F). At the population level, we can reliably decode the identity of the non-task animals in all 3 regions, and consistent with previous results, we only reliably decode task animal identity in the LA after social reward learning (Fig 3G).

Our neuronal recordings also yielded large numbers of neurons from the endopiriform cortex (ENDO). Neuronal activity in the ENDO exhibited properties intermediate to the BLA and PIR, with strong social identity encoding and some social reward learning. These results suggest that the coding of social identity, reward and unexpectedness occurs on a spectrum between the interconnected areas of the rat posterior ventrolateral brain (Fig S6). The brain areas that integrate the most information (BLA and ENDO) also have the highest anatomical connectivity to higher-order cognitive regions^25,26^.

### Representation of conspecifics during natural social exploration reflects the different learning dynamics of the amygdala and piriform cortex

In previous work, we have shown how putative pyramidal neurons in the amygdala were highly tuned to conspecifics and how neurons in the ventral BLA strongly discriminated between male and female rats. To understand how the neuronal encoding of social identity, valence and unfamiliarity seen during the social reward task related to these previous results, we allowed implanted rats to sequentially engage with each of the 4 conspecifics in an open-field environment (Fig 4A) after task learning. We tracked the position of the two interacting rats in the environment to calculate the interindividual distance. Overall, rats did not show any significant differences in the time they spent in close proximity (< 10cm) to the S+, S-, NM and F (Fig 4B), but there were significant changes in the distribution of interindividual distance. Specifically, the implanted animal stayed slightly closer to the S+ and NM and further away from S- and F (Fig 4C). We tracked the activity in a subset of the recorded neurons between the social reward task and natural social behavior session (Fig S7).

**Figure 4.**
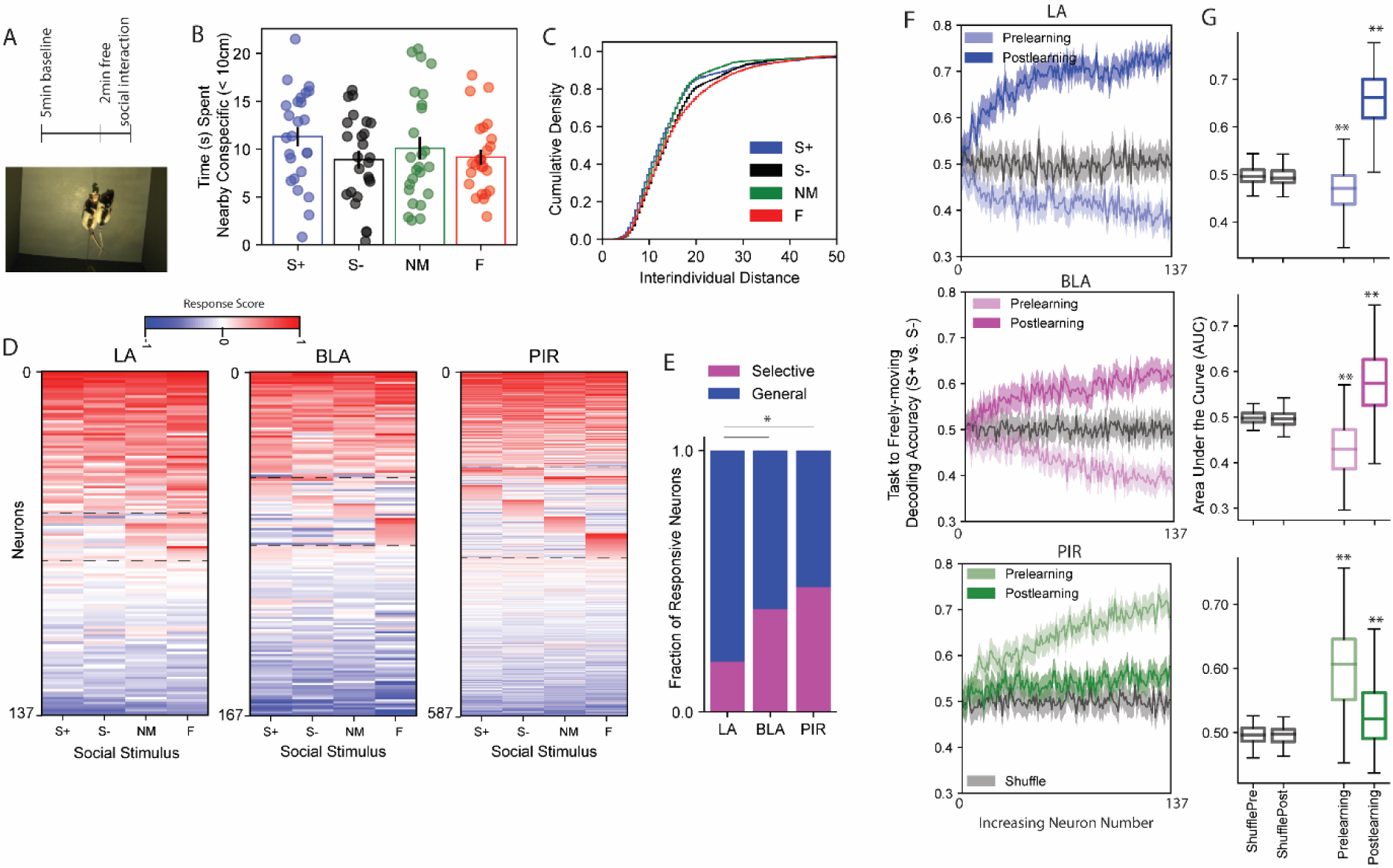
Representation of conspecifics during freely-moving behavior reflects different learning dynamics in the amygdala and piriform. **A)** Conspecifics (S+, S-, NM, and F) were presented in an open-field environment in a pseudo-random manner for ∼2 min each, following a 5min baseline period (top). The implanted rat and stimulus animals were color-coded for dual-animal tracking (bottom). (B) The associated valence of the conspecifics had mild effects on freely-moving social behavior. Rats spent comparable time in direct contact with each conspecific (F(3, 72)=1.65, p=0.19, S+: 11.30±0.99, S-: 8.90±0.91, NM: 10.1±1.16, F: 9.16±0.76, mean±SEM). (C) Overall, rats spent more time further away from the S- and female compared to the S+ and novel male (Kruskal-Wallis test H(3)=884.76, **p<0.01) (D) Single neuron tuning in the LA, BLA and PIR reflected the tuning properties seen during the task. Neurons are divided into 3 categories – broad tuning (top), single-animal tuning (middle) and no selectivity or decreased firing (bottom). The LA and BLA had fewer selective units compared to PIR and this is largely due to an absence in units selectively tuned to the task animals (S+ and S-). (E) Quantification of neuron types revealed differences in selective versus general tuning across regions (LA: 14 / 59; BLA: 33 / 51; PIR: 151 / 165, selective / general). Chi-square comparisons showed significant differences, LA vs. BLA χ2(1, N=157) = 6.6, *p<0.05; LA vs. PIR χ2(1, N=389) = 18.71, **p<0.01, BLA vs. PIR χ2(1, N=400) = 1.60, p=0.21) (F) A SVM classifier was trained using task data from either the pre- or post-learning sensory period with increasing numbers of neurons and tested on the first 20s of natural social behavior. (G) Classifier performance was quantified across 100 iterations by calculating the area under the curve (AUC). Repeated-measures ANOVA with Greenhouse-Geisser correction showed a significant main effect of condition (preshuffle, postshuffle, prelearning, postlearning) in all three regions: LA: F(3, 297) = 445.75, BLA: F(3, 297)=136.72, PIR: F(3, 297)=122.71, **p<0.01. Bonferroni-corrected post-hoc tests revealed significantly higher post-learning decoding accuracy in LA and BLA compared to shuffled and pre-learning conditions (LA: pre = 0.47 ± 0.005, post = 0.66 ± 0.006; BLA: pre = 0.43 ± 0.007, post = 0.58 ± 0.008). In contrast, PIR showed an opposite trend, with higher decoding before learning (pre = 0.61 ± 0.008) than after (post = 0.53 ± 0.005), suggesting stable identity encoding in PIR and learning-dependent enhancement in amygdalar regions.

Single-neuron tuning to the conspecifics during natural social behavior reflected the patterns of activity seen during the social reward task (Fig 4D-E). In the LA, units were broadly tuned to conspecifics, with almost no units selectively tuned to the task animals (S+ and S-) and some units selective for the probe trial animals, while the PIR contained a sizeable population tuned to each of the animals, mirroring the different social identity encoding seen in the task. The BLA once again showed mixed tuning with a preference for probe trial animals.

To test how the neuronal activity during the task sensory period correlated with natural social behavior, we trained a SVM classifier on the population responses during the sensory task period and tested its ability to decode conspecific identity from the first 20s of natural social behavior (Fig 4F). In LA and BLA, only the decoder trained on post-learning neuronal activity accurately decoded conspecific identity (S+ vs. S-) during natural social behavior. In contrast, the pre-learning activity misclassified the conspecific identity (Fig 4G). This suggested that the learning induced changes during the social reward task were stored in amygdala circuitry and reflected in future social interactions. In contrast, in PIR the neuronal dynamics present before learning more strongly decoded conspecific identity during natural social behavior suggesting a transient role for the remapping of conspecific representation in PIR during the previous social reward task.

Using the spike width of the neuronal waveforms, we identified putative pyramidal neurons and interneurons in the three regions (Fig S8A). During social reward learning, interneurons in the three regions played an outsized role in the improvements in decoding accuracy during learning compared to pyramidal neurons (Fig 5), suggesting that the rapid flexible changes in firing activity due to social reward learning occurred primarily in the interneurons. In contrast, pyramidal neurons were more critical for decoding of conspecific identity during natural behavior. This suggests a model where the short-term dynamic changes occurring in the interneuron population may modify the firing activity of pyramidal neurons over time. (Fig S8E-G).

**Fig 5.**
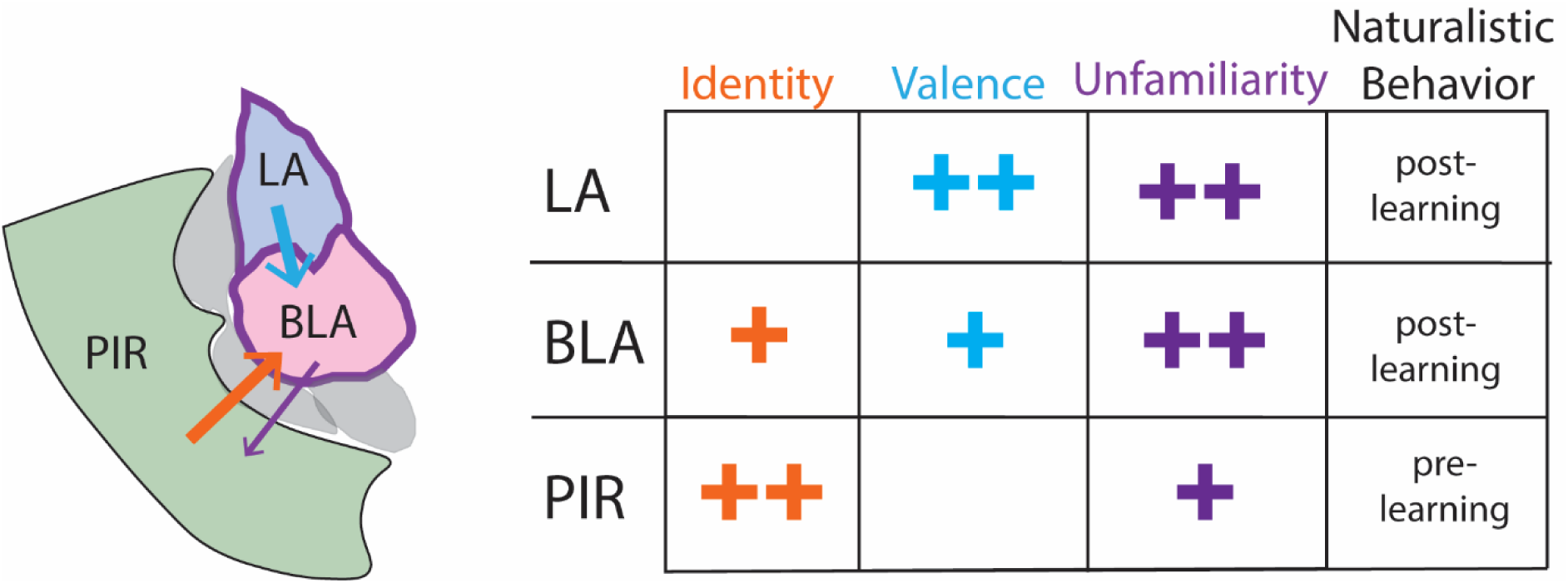
Social identity, reward and unexpectedness in the rat posterior ventrolateral brain. **Lateral Amygdala:** Exhibits little to no social identity encoding for male conspecifics and only with social reward learning do these representations become more distinct. **Piriform Cortex:** Shows distinct representations for all individuals regardless of their reward contingency, task-involvement or sex. **Basolateral Amygdala:** Information about social identity, social reward, and task-involvement is combined in this region, which receives input from both the lateral amygdala and the piriform cortex.

### Non-social odor identity is represented in the LA and PIR

We recorded from a subset of test rats on a non-social odor identity variant of the task (Fig S9A), replacing the stimulus conspecifics with non-social odor extracts (e.g. vanilla, almond). Despite never having experienced a non-social version of the task, rats immediately generalized their task knowledge (Fig S9B-D), which allowed us to compare neuronal responses within the LA, BLA and PIR before and after learning non-social olfactory valence.

We were able to decode the odor identity (O+ vs. O-) from neuronal activity before and after learning in all three regions (Fig S9E-G). Similar to social learning, only the LA and BLA showed significant increases in odor decoding following learning. However, unlike social coding, the LA and BLA had strong decoding of odor identity for the onset of recording. The PIR was able to decode odor identity and this decoding accuracy decreased with learning, suggesting that the PIR does not robustly integrate valence. At the single-neuron level, the LA and PIR had different numbers of neurons responding to the O+ versus O-prior to learning and only the BLA showed a similar reward learning effect to that seen with social conspecifics (Fig SH-J).

## Discussion

### Social identity and valence learning are dissociable in a social reward task

Rats were trained on a social reward task. During a single session, we presented them with multiple conspecifics and measured the acquisition of associative reward learning on a trial-by-trial basis. As in other forms of olfactory-reward association learning^27^, rats improved their accuracy across and within sessions, enabling us to compare learning across animals and recording days.

Interestingly, differences in neural activity emerged before social reward learning, even in sessions where the implanted rat had no prior exposure to the stimulus conspecifics. Almost immediately upon probe trial presentations, rats sampled these animals for longer than the expected task animals. Neurons in the BLA and PIR increasingly encoded social identity or individual recognition, and in the PIR, distinct populations of neurons were recruited for all four animals even when looking at only the first 5 presentation.

Using the social reward task, we were able to measure how neuronal responses to a complex social stimulus dynamically change with learning. Typical social behavior experiments (e.g. 3-chambered sociability test^28^, social memory test^29,30^) rely on measures such as total exploration time to establish recognition and learning. These measures are often slow and highly variable across individuals, which limits their utility in examining the neural correlates of social identity. In contrast, our trial structure allowed us to focus on the social sampling epoch and correlate changes in neural representation of conspecifics with the acquisition of social reward learning.

### Reward learning and stimulus tuning in the LA and BLA

Expanding upon previous research on the amygdala’s role in fear conditioning^31,32^, researchers have described how subsets of neurons in the basolateral amygdala complex (including LA and BLA) exhibit responses to learned appetitive and aversive cues^24,33,34^.

Taken further, these studies suggest that amygdala neurons may encode valence not only in classical conditioning paradigms, but also for innate and learned events in naturalistic situations.

During our social reward task, we observed evidence for valence encoding, however this neural code coexisted alongside strong single-unit tuning to salient conspecifics. Social reward or valence encoding was primarily driven by putative interneurons of the LA and BLA, which developed a rate coding preference or bias to the S+/S-rather than an all-or-nothing on/off firing pattern. These neurons typically responded non-selectively to multiple stimuli and then dynamically increased or decreased their firing activity during learning. In contrast, salient or unexpected conspecifics like the probe animals, strongly activated LA and BLA single-units with all-or-none on/off firing, a pattern observed during both the social reward task and natural social behavior.

The presentation of the probe trial conspecifics during the task could be described as a surprise, a phenomenon distinct from novelty^35^. However, the amygdala responses to these probe trial animals do not align with a simple surprise or arousal signal, as the two probe animals activate distinct populations of neurons. Instead, we hypothesize that these strong responses represent innate stimulus tuning, mirroring the stimulus responses reported in our previous work^13^. The task animals did not recruit the same strong single-neuron responses, which may indicate that the selective amygdala neurons are only recruited in highly arousing or salient situations and that the initial learning of the task structure effectively disengages these strong amygdalar responses.

In the LA and BLA, we observed similar valence-related changes in firing activity, but we identified large differences in single-unit stimulus tuning. The BLA demonstrated stronger social tuning to all 4 social conspecifics especially between different male conspecifics, while the LA exhibited less selective social tuning among same-sex conspecifics but remarkably strong tuning to non-social olfactory odors. Together with recent studies^13,36^, this supports a spatial localization of stimulus responses within the broader amygdala in part due to different input/output connectivity between amygdala sub-divisions^37^.

Taken together, this research proposes a way of unifying previously distinct lines of amygdala research. Neurons of the BLA complex display different modes of neuronal coding, with a modulation of firing rate or dynamic code that signals valence^38^ and recruitment of single-units exhibiting all-or-none firing to stimuli like the probe trial animals. Highly salient events like fear conditioning or food deprivation may change the balance of single-unit responses. Future research is needed to understand when and how the interaction between these neural codes occurs, and to link these neuronal codes to the amygdala’s role in action selection^39^, state^17^, concept-encoding^40^ and memory consolidation^41^.

### Social identity coding and remapping in the PIR

The piriform cortex exhibited the strongest social identity encoding of all 3 areas. One interpretation is that the unique olfactory mixtures that make up social odors for each conspecific activated a unique ensemble of PIR neurons, similar to how the PIR responds to non-social odor mixtures and single-molecule odors^5,42^ . Previous studies have shown that the representation of sensory responses in olfactory and visual cortices were not stable^43–45^.

We also find that a large fraction of PIR responses to conspecifics remap or drift during the social reward task session, which lasted 1-2 hours on average. This remapping of sensory representation did not improve decoding accuracy, consistent with prior studies showing that reward modulation of sensory stimuli in the PIR is minimal, especially in comparison to other regions of the olfactory system^46^.

Recently, more attention has been drawn to studies showing that sensory cortices encode other contextual factors apart from their unique sensory representation^6,7^. We also find that the piriform cortex encodes the probe animals differently from the task animals but using different coding principles from the amygdala. Specifically, the less selective or general neurons in the PIR responded more strongly to the probe animals, though overall the PIR did not have as strong a response as the amygdala. One possibility is that the information about the unfamiliar conspecific probe animals is communicated from an area like the amygdala to the PIR.

### Understanding task responses in the context of natural social behavior

Neuronal activity during natural social interaction can be difficult to interpret compared to tasks with repetitive, well-defined behavior. However, an understanding of how neural activity during a controlled task correlates to real-world behavior is critical for fully understanding how the brain works^47^. In the three brain regions studied here, we see similarities in neuronal responses between the task and natural behavior with increased identity coding from the LA to the PIR particularly in neurons selectively tuned to individuals. Interestingly, we find key differences in how the task responses correlate to natural social behavior. In the amygdala, the post-learning task activity best reflected the neuronal dynamics during natural social behavior. In contrast, it is the pre-learning PIR activity which best represents the naturalistic behavior. This suggests that the information about social identity and reward are stored in different ways in these two structures. In the amygdala, learning-induced plasticity is preserved in the circuitry and then reactivated in a different context. In the PIR, the remapping of activity during the learning task is transient and less relevant to social behavior than the initial social identity representation.

In conclusion, we find differential roles for the LA, BLA and PIR in encoding social identity, reward and unfamiliarity of social conspecifics (Fig 5). Social identity information is strongest in the olfactory sensory piriform cortex while social valence is strongest in the LA. These information streams along with strong signals for unfamiliar conspecifics converge in the BLA which acts as a node connecting this sensory information with higher-order brain areas like the prefrontal cortex and hippocampus to influence decision-making and memory.

## Supporting information

All Supplemental

## Acknowledgements

We thank the SWC FabLabs especially Simon Townsend for their support and assistance in manufacturing and building the rat social reward task. We thank Tianyi Zeng, Laura Schwarz and Margaret Conde Paredes for their technical support and to members of the O’Keefe Lab for helpful discussions. This project was supported by the Kavli Foundation (LS-2022-GR-25 awarded to CM), the Sainsbury Wellcome Centre Core Grant from the Gatsby Charitable Foundation and Wellcome trust (090843/F/09/Z) and a Wellcome Trust Principal Research Fellowship (Wt203020/z/16/z awarded to JO’K).

## MATERIALS AND METHODS

### Animals

Rats used in these experiments were adult male or female Lister Hooded rats obtained from Charles River, UK. 20 male rats were trained in the social reward task, and 10 of these rats received chronic Neuropixel implants. 44 male rats and 4 female rats were used as stimulus conspecifics. All rats were aged between 2-6 months at the start of the experiment, and animals receiving Neuropixels implants weighed 350-450g at the time of surgery. Rats were housed in pairs in large clear polycarbonate cages, except for post-surgical rats, which were housed individually. Cages were located in a temperature- and humidity-controlled room with a 12-hour reverse light/dark cycle. All experiments were run in the dark phase. All rats received ad libitum access to water. Conspecific rats also had ad libitum access to food. The 20 rats trained on the social reward task were fed once per day and maintained at least 90% of their pre-restriction body weight. All animal procedures were performed in compliance with the UK Animals (Scientific Procedures) Act (1986).

### Automated Social Reward Training Chamber

Training took place in a custom-made automated operant chamber (40×40×40 cm) equipped with a sensory sampling port and a food port on opposite sides. Conspecifics were presented in boxes mounted on an adjacent rotating platform driven by a servomotor (Dynamixel XM430-W350-T) connected to a gear system. An opaque sliding door, operated by a linear actuator and linear actuator control board (P1-P Linear Actuator, P16-100-22-12-P, Actuonix) separated the operant chamber and the rotating platform. Fans (668-8839, RS Pro) were located on top of the stimulus rat boxes and on the sides of the operant chamber (412 series, ebm-papst). A pellet dispenser (80209, Campden Instruments) dispensed 45mg food pellets (sucrose or chocolate grain-based precision pellets, Bio-Serv) into the food port. A small LED (8mm, 600Mcd, white, Kingbright) located in the operant chamber signaled trial availability.

Rats’ location and actions were detected using several sensors. A light curtain with an optical crosshatch (emitter and receiver, LX3ESR and LX3RSR, Turck Banner Ltd) detected presence at the sensory sampling port. A pair of infrared sensors (3mm LEDs, ID 2166, Adafruit) in the food port detected nose pokes. An infrared sensor (Omron EE-SX3160-W11) monitored the opening of the sliding door for conspecific presentation.

Two video cameras (Basler acA2500-60uc) monitored the stimulus and test rats. All sensors and motors were wired to 2 connected Arduinos (Arduino Mega 2560 Rev3), which were controlled by a custom-written Matlab program. Digital signals for all the sensors, the LED signalling trial availability and the motors were wired into and recorded using an Intan RHD USB Interface Board at 1kHz sampling rate. For Neuropixels recorded, the digital signals were wired into a National Instruments board (SCB-68A Noise Rejecting, Shielded I/O Connector Block) and saved alongside neural signals.

## Experimental design

### Initial Training

We trained adult male rats in a Go/No-Go social reward task inspired by a previously published social discrimination task^23^. Prior to being trained on the social reward task, rats were pre-trained in four phases, typically completed across four days.

Phase 1 (60 trials): Rats automatically received a food pellet at the food port after poking the sensory sampling port for 100ms.

Phase 2 (60 trials): Sensory sampling duration required for reward increased incrementally from 100ms to 1000ms in 250ms steps.

Phase 3 (60 trials): A single stimulus rat (S+) was introduced. To successfully complete a trial during this phase, the test rat was required to initiate a trial (100ms poke at sensory port) which opened the sliding door. It then needed to sample the stimulus conspecific for at least 1000ms before a pellet was dispensed at the food port on the other side of the box.

Phase 4 (60 trials): Pellets were no longer automatically dispensed after sensory sampling. Rats had to poke at the food port within 10s after sampling the stimulus rat. If the test rat failed to nose-poke the food port within 10s, the trial was counted as a miss and no reward given. Performance of >80% was required to progress to social discrimination training.

Phase 5 (social reward task): After completing pre-training, rats were trained on the social discrimination task during daily sessions of 120 trials. Stimulus rats were 2 age-matched male rats housed in different cages in the same holding room. Prior to training, they were assigned as rewarded (S+) or unrewarded (S-) and this valence was maintained throughout behavioral training. Both test and stimulus rats could move freely in their respective compartments during training. Each stimulus rat was placed in a different box of the rotating platform on successive days. Between sessions, all parts of the setup were cleaned with 70% ethanol.

At the start of each trial, the LED near the sampling port was activated and all the fans powered on. The test rat initiated the trial by poking the sensory sampling port for 100ms, activating the sliding door to present one of the two conspecifics. Test rats were required to sample the conspecific for a minimum continuous period of 1000ms with no upper time limit. Once the minimum sampling time was met, withdrawal from the sampling port started a counter for the 5s response window and closed the sliding door. If the S+ was presented, a nose-poke at the food port during the response window activated the pellet dispenser and counted as a hit, whereas no nose-poke at the food port counted as a miss. If the S-was presented, a nose-poke at the food port during the response window counted as a false alarm which led to a 15s delay in the start of the next trial, whereas no nose-poke counted as a correct rejection. After the sliding door had closed and the response window plus any delay had passed, the revolving platform rotated into place for the next trial. The inter-trial interval (ITI) was 15s.

In each session, there was a total of 120 trials with 60 presentations of each conspecific in a pseudo-randomized fashion. In a single block of 20 trials, the number of presentations of each stimulus animal was balanced as was the direction of clockwise or counter-clockwise rotation. Each stimulus animal was presented no more than 3 times in a row. During initial training, a performance of ≥ 75% correct responses over all trails during a daily session was defined as criterion. Out of the 20 rats, 17 learned the task and reached criterion. 3 rats failed to learn the task within 15 days and were excluded from the study.

### Generalization

After reaching criterion, a subset of rats (N= 10) was further trained on a generalization version of the task. A new pair of male conspecifics (age-matched, housed in different cages in the holding room and having no prior exposure to the test rat) was presented, and the criterion set to 80% correct in a 20-trial block. Rats were trained for 120 to 360 trials per day, and training stopped after criterion was reached. After initial training, generalization training was repeated with a minimum of 2 additional pairs. The last pair learned during generalization training was the first pair presented after Neuropixels implantation.

### Post-implant Behavior Schedule

After successful generalization training, rats underwent surgery to chronically implant Neuropixels probes to the left posterior ventrolateral brain. After 48h recovery, rats performed 2 sessions on separate days of the probe-trial variant of the social discrimination task. Rats completed 120 trials to 360 trials or until they lost motivation. After each session rats engaged in natural social behavior with each of the four conspecifics in an open-field environment. If good quality electrophysiological recordings were still possible after the 2 sessions of social reward learning, rats performed the non-social odor reward task.

Experiments were repeated if additional neuronal activity was present on other electrodes of the Neuropixels probe.

### Probe-trial variant of social reward task

During Neuropixels recordings, we presented either a novel male or female conspecific as probes during the typical task structure. These probe trials were uncued and occurred after every block of 10 trials. The trials were not rewarded but also did not cause a timeout or task delay. The novel male conspecific was an age-matched male conspecific that had had no prior exposure to the implanted animal. The female conspecific was an adult female rat.

During post Neuropixels implantation recording, the novel male conspecific was changed every day, while the female remained the same throughout.

### Non-social odor reward task

For 2-odorant reward learning, rats were presented with odorants instead of conspecifics. Rats had no experience with any of the odorants before the experiment. 1 ml of undiluted odorant was transferred into a weigh boat and put inside the conspecific boxes. The following odors were used: Orange blossom (Nielsen-Massey), Sicilian Lemon (Dr. Oetker), Almond Extract (Dr. Oetker) and Vanilla extract (Nielsen-Massey).

### Naturalistic Behavior in the Open-Field

After successful social reward learning, implanted rats were returned to their home cages for 1-2h and given food. Rats (implanted and conspecifics) were lightly colored to facilitate multi-animal tracking (see ^13^) and implanted rats were then reconnected to the Neuropixels cables in a separate environment (100x70x40cm LxWxH, black acrylic box) illuminated by 2 LED photo lights (F&V K480 Lumic Camera Lamp, Wex lighting). Rats were allowed to freely explore the environment for 10min before conspecific introduction. Each of the conspecifics was introduced into the environment where the implanted rat was for 1.5-3min and a 5min baseline recording separated each of the conspecific presentations. Rats were allowed to freely-interact and experimenter-intervention only occurred when rats were aggressive or there appeared to be danger to the implant. The setup was lightly cleaned with 70% ethanol between presentations. Conspecifics were presented in a pseudo-randomized fashion where the female conspecific was always presented last. Presentation order of the S+ and S-so that the S+/S-were presented first in 50% of the recording sessions for any single rat.

### Olfactory and Visual Block Experiments

A subset of 5 rats that had undergone the initial training phase (but not generalization or Neuropixels implant) were challenged by blocking a specific sensory cue during the social reward task. We blocked olfactory input by removing the holes that allowed passage of air flow from the conspecific boxes to the operant chamber and disconnecting the fans located on top of the conspecific boxes. We blocked visual input by exchanging the clear acrylic sheet separating the test rat and the stimulus rats for an opaque acrylic sheet with a single air hole. We counter-balanced which sensory cue was blocked first (i.e. half the rats received olfactory block first and the other half visual block first). Most rats recovered performance after the sensory block was reversed the following day. These rats were not used for Neuropixels implants or further experiments after sensory block.

### Neuropixels implantation

To perform electrophysiological recordings, rats underwent stereotactic surgery to implant a single 4-shank Neuropixels 2.0 probe^48^ targeting the left posterior ventrolateral brain, including BLA, LA and posterior PIR. To protect the implant, we used a modified 3D printed head cap system (based on ^49^). We used both Repix holders^50^ and metal microdrives^49^ to attach the probes to facilitate later probe retrieval. Rats received pre-operative subcutaneous injections of dexamethasone (0.7mg/kg) to improve neuronal yield and stability^50^. Rats were anaesthetized with isoflurane in O2 (2-3%) and given pre-op analgesia subcutaneously (Carprieve 5mg/kg) before being placed in a stereotaxic device. All 10 rats were implanted in the left hemisphere (coordinates relative to bregma, 3.2-2.8mm posterior, 4.7-5mm lateral and -9mm ventral).

Prior to implantation, Neuropixels probes were dipped in 1 µL of CM-DiI solution (Thermo Fisher, 50 μg dissolved in 50 μL of pure ethanol ^51^). After exposing the skull, we fixed the head cap base, a stainless-steel reference screw and additional support screws to the skull using Super-Bond (Sun Medical) and light curable epoxy (RelyX, 3M). After craniotomy and durotomy, we slowly inserted the Neuropixels probe to maximal depth to target deep-brain regions using a micromanipulator (SM10 compact, Luigs & Neumann). The base of the probe holder was cemented in place and the probe detached from the stereotax holder. The exposed craniotomy was covered with a thin layer of Duragel (Cambridge Neurotech) and sterilized Vaseline. The ground screw was wired to the probe and the Neuropixels headstage was secured onto the head cap. The opening of the head cap was wrapped with 3M Coban wrap when the probe was not in use.

Post-surgery rats received analgesia and antibiotic mixed in strawberry jelly for 3 and 5 days respectively (Metacam 1mg/kg, Baytril 1%) and sub-cutaneous injections of dexamethasone at day2 and 5 post-surgery (0.2 mg/kg).

Neuropixels probes were retrieved for reuse after the experiment was completed. Briefly, the rat was anesthetized as above and the head cap removed. Probes were unscrewed from their cemented base and removed, tested, cleaned (1h in 1% Terg-a-zyme in MilliQ, Sigma-Aldrich, followed by 1h in milliQ) and in some cases reused. Anaesthetized rats immediately received an overdose of sodium pentobarbital prior to perfusion.

### Histology

After the administration of sodium pentobarbital, rats were transcardially perfused with phosphate-buffered saline (PBS) and 4% paraformaledyde (PFA). After at least 24h in PFA, brains were transferred to phosphate buffer (50mM PB) and then coronally sectioned in 50um steps while simultaneously imaged using a custom serial two-photon microscope.

There is currently no available rat brain atlas with defined borders for the BLA for automatic registration, therefore location of each of the 4 shanks of the probe were determined by manually registering the Di-I tract onto the Paxinos brain atlas and mapping the coordinate channel locations of the Neuropixels probe onto the tract.

### Data collection/experimental setup

Neuropixels recordings were collected as described in ^50^. Briefly, Neuropixels were connected via cables to an IMEC card housed in a National Instruments chassis (NI PXIe-1071) and data were acquired at 30KHz using SpikeGLX software (https://github.com/billkarsh/SpikeGLX). Task and camera timestamps were recorded alongside the electrical signals in SpikeGLX and cameras were controlled using custom-written Bonsai scripts^52^ during the task and custom-written Labview scripts (National Instruments with NI USB-6000) during naturalistic behavior. During task learning, neural data was transmitted through an electrophysiology commutator (Assisted Electrical Rotary Joint, Doric Lenses) to neutralise the turning of the rat.

48h after surgery, rats were connected for an initial baseline recording and probe survey. Given the electrical constraints on the 4-shank Neuropixels probes, we were restricted to recording from 384 channels out of a possible 5120 at any given moment. We selected an initial channel map and completed a full set of behavioral experiments (2 social reward sessions, 1 non-social olfactory session). At the conclusion of these experiments if viable electrophysiological activity was still present on channels not previously recorded from, we selected a second channel-map and conducted another round of experiments.

### Single-cell isolation from multi-unit recordings

Neuropixels data was concatenated across a session (social reward task and naturalistic behavior were kept separate) and processed and filtered using CatGT (https://billkarsh.github.io/SpikeGLX) and Kilosort 2.5. The data was manually curated using Phy2.0 (https://github.com/cortex-lab/phy) according to the following criteria: less than 0.1% of spikes violated the cell refractory period of 2ms, spike waveform consistent with single unit activity, amplitude of at least 40mV and absence of any 50Hz noise in autocorrelogram. Waveform and cross-correlograms of all nearby units were compared to verify that there were no two clusters corresponding to the same neuron.

### Tracking Single-units across recording sessions

To identify single-units that were continuously present during the social reward task and the subsequent naturalistic behavior, we combined existing tools^53^ with our prior methods (structural similarity index, SSIM used in ^13^). Using custom-written python code we overlay normalized waveforms extracted from high-density Neuropixels and visually verified that each identified unit was matched to a single other unit present in the other recording session. A larger percentage of units was successfully tracked in lower-density areas like the amygdala compared to higher-density recording areas in the piriform cortex.

### Classification of putative pyramidal neurons and putative interneurons

We excluded neurons with inverted waveforms (where the peak voltage precedes the trough) and calculated the time delay between the trough and the peak voltage for the largest amplitude neuronal waveform from each neuron. Since we were comparing waveforms across 3 areas with neurons with distinct morphological features, we used a simple spike width cut-off of 500us to divide the population into putative interneurons and putative pyramidal neurons.

### Tracking from video data during naturalistic behavior

We tracked the position of the two rats in the open-field chamber (for full details see ^13^). Briefly, rats were lightly colored prior to the experiment to facilitate tracking. Video data was processed offline to identify the coordinate positions (DeepLabCut 2.3.8^54^) and we calculated inter-individual distance between the implanted rat and the conspecifics.

### Behavioral Learning

Per-trial performance was calculated by taking the per-trial accuracy and calculating the rolling mean over 20 trials. We divided sessions into pre- and post- learning trials by calculating the learning inflection point, defined as the time when performance reliably crosses threshold (70%). To ensure we looked at sensory sampling tied to pre- or post- learning performance, we included only hits and false alarms in pre-learning data and only hits and correct rejections in post-learning data. On rare occasions, rats lost motivation at the end of the recordings (as defined by continuous misses and correct rejections), and we excluded all trials during these times.

We identified specific times in each trial that signaled sensory input and decision-making and used these to analyze neural activity. The onset of the sensory period was defined by the activation of the door sensor allowing interaction between the implanted and stimulus rat.

The end of the sensory period was defined as the time when the rat initiated the 5s response window countdown or 3s after the start of the sensory period, whichever came first. The action period started from the initiation of the response window countdown to the time of nose poking the food port or the end of the response window, whichever came first.

## Analysis

### Comparing neuronal activity across trials of different lengths

The firing rate of each neuron across each individual trial was time-warped to fit a common time frame. Each behavioral trial was divided into 4 epochs (baseline, sensory, action and outcome) to compare neuronal firing data between trials of different durations. The start and stop of the sensory and action epochs is based on the sensors in the behavioral apparatus as previously defined. Baseline firing is the 5s of activity preceding sensory start and outcome is the 5s of activity following the end of the action epoch. Neuronal firing rate was calculated using 100ms bins and time-warped into 40 bins (10 for each epoch) using linear interpolation (scipy.interpolate.interp1d).

### Classifying task responsiveness of individual neurons

We evaluated the responsiveness of each individual neuron during the sensory period during pre- and post- learning conditions. We calculated the mean response of each individual neuron to the rewarded and unrewarded stimuli before and after learning and compared the firing activity during the baseline period to the sensory period using Wilcoxon rank-sum test. All individual neurons with a p value less than 0.05 were considered responsive for a given condition (4 conditions, S+ pre-learning, S- pre-learning, S+ post-learning, S- post-learning). When comparing responses among only responsive neurons, we included all neurons that were significantly responsive for at least one condition.

### Decoding of stimulus identity

To test whether stimulus identity could be reliably decoded from population activity under different conditions, we used a support vector machine classifier (SVC) with a linear kernel function (SVM, python package sci-kit learn). We divided data into training/testing datasets using 5-fold cross-validation and standardized the neural data (sci-kit learn StandardScaler) before running SVC classification. Results were calculated from neuronal data from the same brain region but across multiple rats. All datasets were balanced in the number of trials included (S+, S-, pre- and post-learning) and the number of neurons used to calculate results. Overall results shown are the mean +/- sem of 100 iterations to ensure that all neurons and all trials are ultimately represented.

To calculate decoding accuracy across the trial, we used the time-warped neuronal data and calculated the decoding accuracy for each time bin. To look at the effect of increasing numbers of neurons on stimulus decoding, we calculated the decoding accuracy during the sensory period with increasing neuron counts from 2 to 348 neurons with 100 iterations for each neuron count. For 4-animal decoding conditions, we included probe trials in both the pre- and post-learning datasets and reran the SVC classification and calculated the confusion matrix (confusion_matrix, sci-kit learn). To test whether pre-learning data can decode stimulus accuracy in the post-learning data and vice versa, we selected a balanced pre- and post-learning dataset based on the minimum number of available trials and trained the SVC classifier on all the selected trials of the pre-learning dataset and tested on all selected post-learning trials. This was repeated across 100 different iterations.

To compare decoding accuracy from the task to naturalistic behavior, we reduced task responses to a single value response score by calculating the receiver operating curve (ROC) for each neuron comparing baseline to sensory firing activity for either pre- or post-learning data and quantifying the difference by calculating the area under the curve for each neuron and each trial. We then calculated the response scores during naturalistic social interaction using a sliding window of 5s across the first 20s of social interaction. Each neuron used in this analysis was previously identified as a continuous single-unit and had response score values for the social reward task and naturalistic social behavior. We trained an SVC classifier on the task responses and tested it on the activity during naturalistic behavior. The accuracy is the mean accuracy from the first 20s of social interaction.

### Population correlation of responsive neurons

To calculate the correlation between the population across different conditions (S+, S-, pre- and post-learning), we calculated the per-unit sensory response score by quantifying the receiver operating curve response during the sensory epoch. We calculate the Pearson’s correlation between the population vectors comprised of the sensory response score for each responsive unit in a given condition (e.g. S+ pre-learning).

To calculate the correlation between the first 5 presentations of each of the 4 conspecifics (S+, S-, NM and F), we calculated the average firing rate per neuron during the sensory epoch and calculated the Pearson’s correlation between the resulting population vectors.

### Neuronal trajectories and trajectory distance

To calculate the neuronal trajectory during the sensory period for each of the 4 conspecifics presented, we z-scored the mean activity per neuron for each stimulus presentation. Data from 50 randomly selected units were concatenated and dimensionality reduced using principal component analysis. This was repeated 100 times selecting a different subset of 50 neurons and each brain area was calculated separately. The mean trace of the first three principal components was plotted for visualization. To quantify the trajectory distances, we calculated the Euclidean distance of the neural trajectories during middle of the sensory epoch for each conspecific and every iteration.

### Example Raster Plots and PSTH

Raster plots and PSTHs for example cells were made using the MLIB library.^55^

