## Supplementary material for "Differential encoding of social identity, valence and unfamiliarity in the amygdala and piriform cortex": All Supplemental

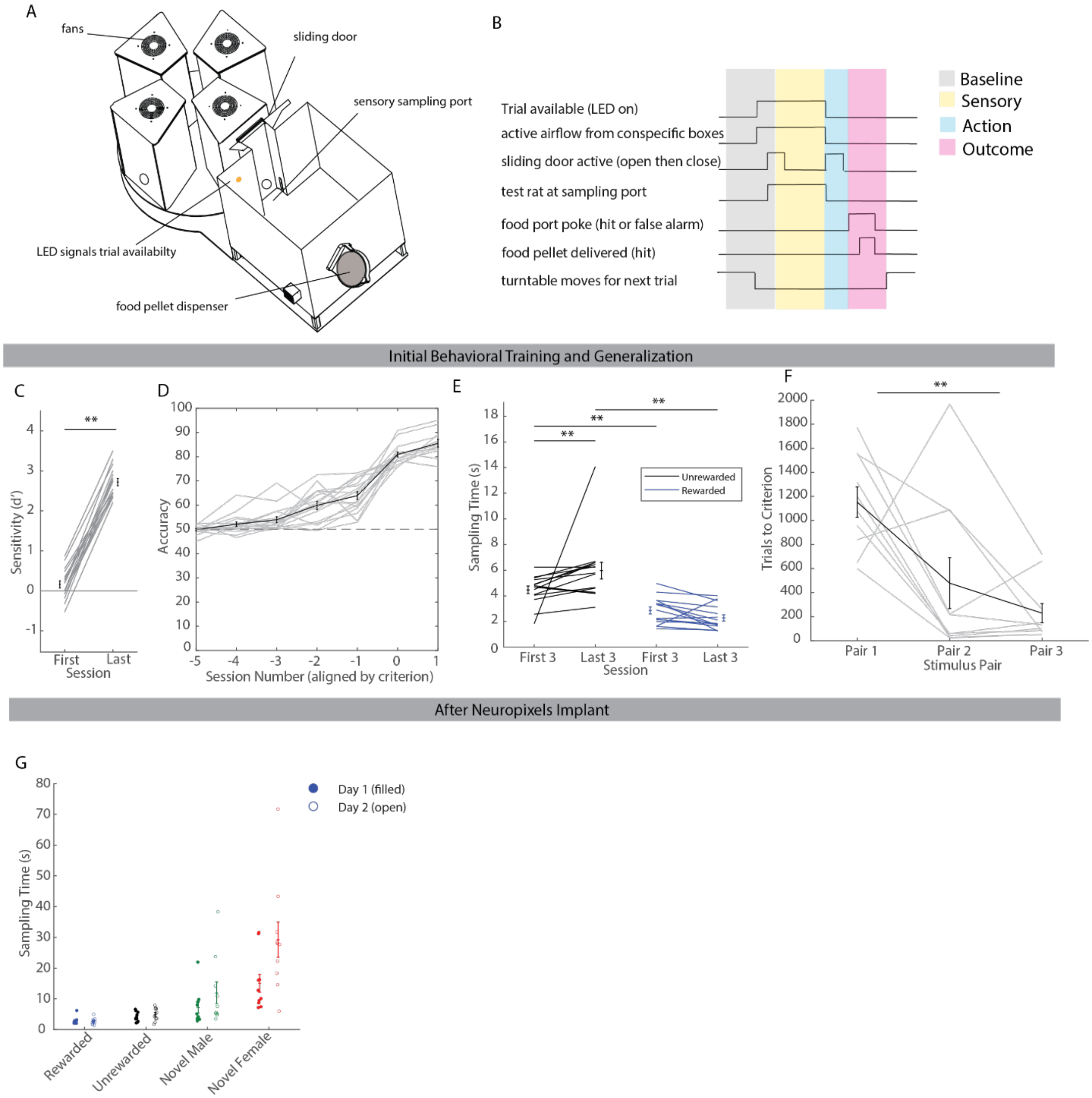


**Figure S1. Social reward task learning and performance after surgery**

**(A)** A more detailed schematic of the behavioral setup.

**(B)** An overview of the temporal order of events during a trial. When a new trial is available, an LED light in the test chamber is activated and the fans above the conspecific boxes are powered on. Once the test rat breaks the infrared beam at the sensory sampling port, the sliding door opens and the sensory period starts (yellow shaded). The test rat must sample for a minimum of 1 continuous second before making a decision. Once the rat moves away from the sensory sampling port after the minimum sampling time, the sliding door closes and an action/response period (blue shaded) starts. The rat has a maximum of 5s to poke the food port. A nose poke of the food port during the action/response period leads to the delivery of a food pellet in rewarded trials and a 15s timeout in unrewarded trials (outcome epoch – magenta shaded).

**C**) Rats increase their behavioral sensitivity (d') from the first to the last session. First (mean 0.15 +- 0.09 SEM) vs. last session (mean 2.72 +- 0.09 SEM), paired t-test, $p < 0.005$, N=17 rats.

**D**) Accuracy for each rat is aligned to the day of reached criterion. The average difference in accuracy between reaching criterion and the day before is 16.98 +- 1.98 SEM.

**E**) The average sampling times for rewarded and unrewarded conspecifics is shown for the first 3 and last 3 sessions of the initial training phase. A mixed repeated-measures ANOVA was conducted with Trial (First vs. Last) as a within-subjects factor and Reward Status (Rewarded vs. Unrewarded) as a between-subjects factor to examine their effects on sampling time. The main effect of Trial was not significant, $F(1,28) = 1.189$, $p = 0.285$. The interaction effect (Trial × Reward Status) was significant, $F(1,28) = 6.056$, $p = 0.020$. Post-hoc Comparisons (Bonferroni-corrected): First Unrewarded vs. Last Unrewarded: $p = 0.0181$ (not significant after correction), First Rewarded vs. Last Rewarded: $p = 0.3408$. First Unrewarded vs. First Rewarded: $p = 0.0004$. Last Unrewarded vs. Last Rewarded: $p < 0.001$. N= 15 rats (where timestamps were available).

**F**) The number of trials to criterion is shown for the first 3 pairs of conspecifics (mean + SEM). Pair 1 corresponds to the initial training phase. For pair 1, criterion is 75% correct within one session, for the following pairs it is 80% in a block of 20 trials. The trials to criterion differed significantly across the three stimulus pairs ($F(2, 22) = 10.776$, $p > 0.0001$, repeated measures ANOVA). N =10 rats (all implanted rats).

**G)** Following surgery, rats had similar sampling times on day 1 and day 2 for the first 10 trials across the 4 trial conditions. A two-way repeated-measures ANOVA was conducted to examine the effects of: condition (rewarded, unrewarded, novel male, novel female) and day (day1, day2) on sampling time. Both factors were within-subject. There was a significant main effect of condition: $F(3) = 24.36$, $p = 0.000$, but not on day: $F(1) = 4.99$, $p = 0.052$. The interaction condition × day was significant: $F(3) = 4.75$, $p = 0.042$. N=10 rats

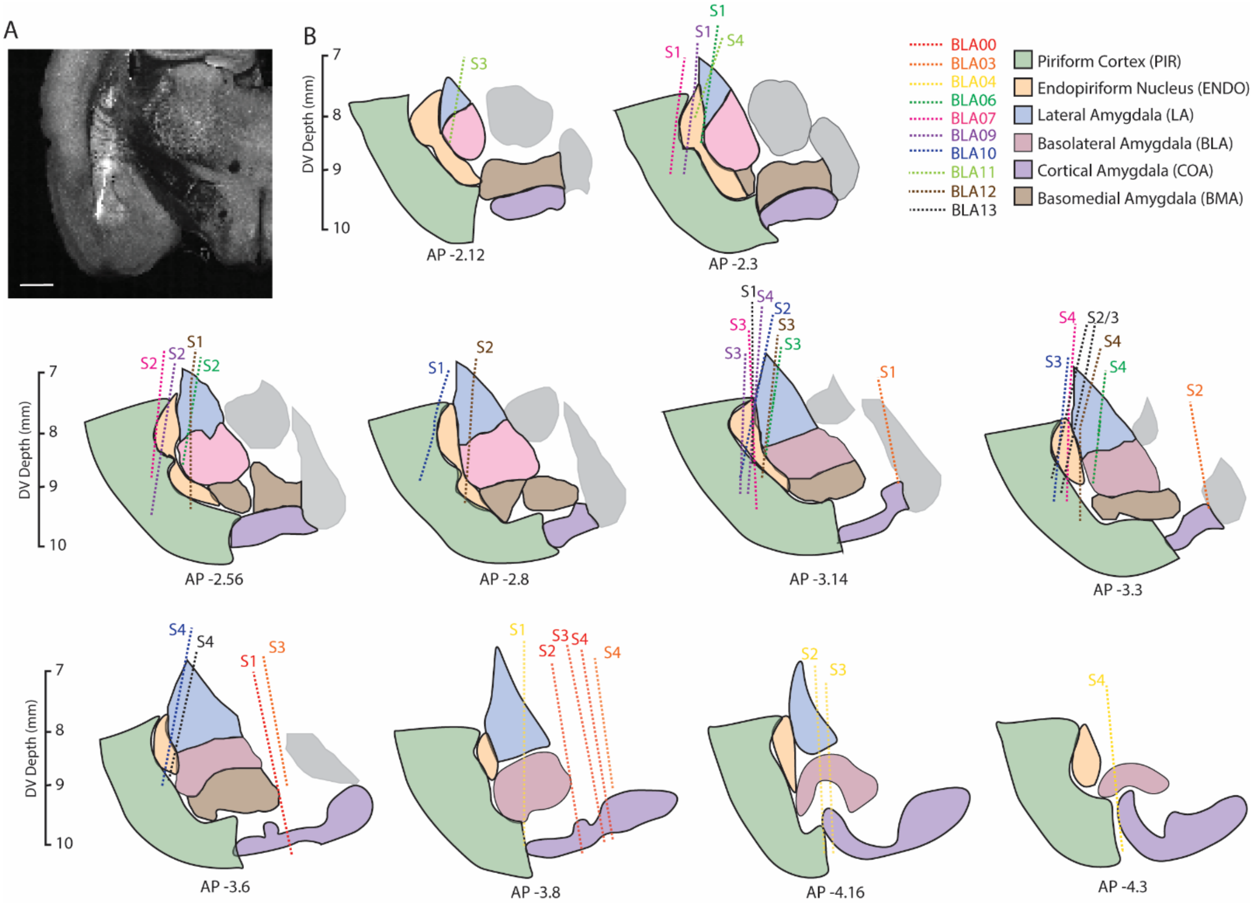


**Figure S2. Targeting to multi-shank Neuropixels probes to the rat posterior ventrolateral brain**

**(A)** A coronal brain slice showing the rat posterior ventrolateral brain and the positioning of 1 of the 4 shanks of the chronic Neuropixels 2.0 implants. The probe is dipped in CM Di-I prior to surgery and post-fixation the brain is sectioned and imaged using 2-photon serial tomography. (scale bar = 1mm)

**(B)** The location of each of the 4 shanks for the 10 implanted rats. Each shank is denoted S1-S4 and color-coded based on the rat identity.

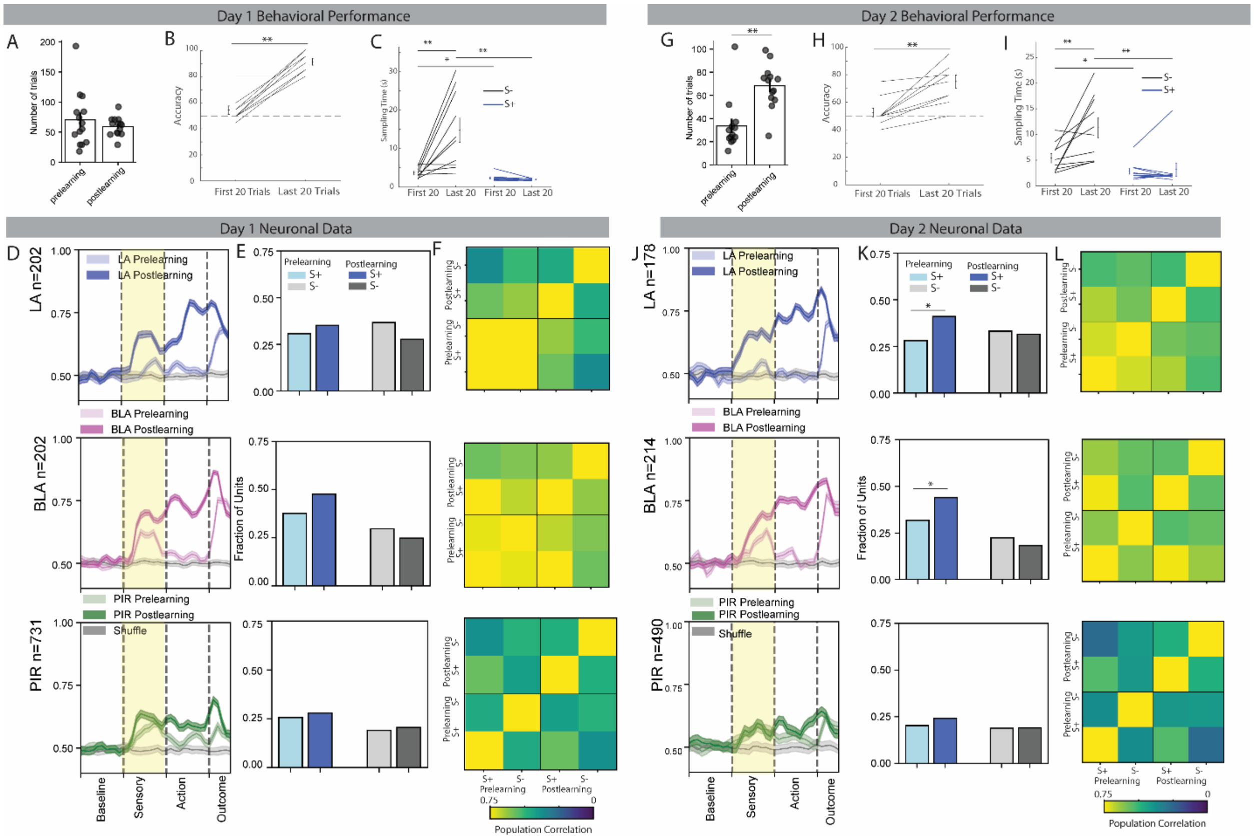


**Figure S3. Behavioral performance and neural recordings were comparable across 2 days of the social reward task.**

**A)** The number of trials that occurred before and after social reward learning on day1 of task performance. Each point represents an individual session from one rat. (p=0.47, paired t-test N=14 sessions).

**B)** Behavioral accuracy increased from the first to last 20 trials on day1 of the social reward task (first session after surgery). First 20 trials (mean 54.5 +- 3.1 SEM) vs. last 20 trials (mean 91 +- 2.56 SEM), paired t-test, $p < 0.001$, N=10 rats.

**C)** Average Sampling times are shown for the first and last 20 trials (Unrewarded: First 3.68 +- 0.4 SEM, Last 14.77 +-3.14, Rewarded: First 2.37 +-0.28 SEM, Last 1.93 +- 0.08 SEM). A mixed repeated-measures ANOVA was conducted with Trial (Early vs. Late) as a within-subjects factor and Reward status (Rewarded vs. Unrewarded) as a between-subjects factor. The main effect of Trial was significant, $F(1,18) = 11.84$, $p = 0.0029$. The interaction effect (Trial × Reward Status) was significant, $F(1,18) = 13.86$, $p = 0.0016$. Post-hoc comparisons (Bonferroni-corrected): Early Unrewarded vs. Late Unrewarded: $p < 0.001$, Early Rewarded vs. Late Rewarded: $p = 0.8446$. Early Unrewarded vs. Early Rewarded: $p = 0.0161$ (not significant after correction). Late Unrewarded vs. Late Rewarded: $p = 0.0007$. N = 10 rats.

**D)** Decoding of conspecific identity (S+ vs. S-) from population neural activity in the 3 regions (LA: top, BLA: middle and PIR: bottom) using a SVM classifier on the first day of social reward training.

**E)** The fraction of units with significant responses during the sensory period changes in response to learning on day1. (change in S+ responsive: LA = $\chi2(1, N=202) = 0.72$, $p=0.40$; BLA = $\chi2(1, N=202) = 3.65$, $p=0.06$; PIR = $\chi2(1, N=731) = 0.89$, $p=0.34$, change in S- responsive: LA = $\chi2(1, N=202) = 3.28$, $p=0.07$; BLA = $\chi2(1, N=202) = 1.01$, $p=0.31$; PIR = $\chi2(1, N=731) = 0.43$, $p=0.51$)

**F)** Correlation of the pre- and post-learning population activity during the sensory sampling period on day1.

**G)** The number of trials that occurred before and after social reward learning on day2 of task performance. Each point represents an individual session from one rat. (**$p<0.01$, paired t-test N=13 sessions).

**H)** Behavioral Accuracy of rats in the second session after surgery (day 2). First 20 trials (52.5 +- 3.18 SEM), Last 20 trials 75 +- 4.71 SEM), paired t-test, $p=0.001$.

**I**) Average sampling times for the first and the last 20 trials (Unrewarded: First 5.39 +-0.90, Last 11.31 +-1.96, Rewarded: First 2.84 +- 0.56, Last 3.18 +- 1.28). A mixed repeated-measures ANOVA was conducted with Trial (Early vs. Late) as a within-subjects factor and Reward Status (Rewarded vs. Unrewarded) as a between-subjects factor. The main effect of Trial was significant, $F(1,18) = 8.42$, $p = 0.0095$. The Interaction effect (Trial × Reward Status) was significant, $F(1,18) = 6.72$, $p = 0.0184$. Post-hoc comparisons (Bonferroni-corrected): Early Unrewarded vs. Late Unrewarded: $p = 0.0011$, Early Rewarded vs. Late Rewarded: $p = 0.8291$, Early Unrewarded vs. Early Rewarded: $p = 0.0284$ (not significant after correction), Late Unrewarded vs. Late Rewarded: $p = 0.0028$.

**J)** Decoding of conspecific identity (S+ vs. S-) from population neural activity in the 3 regions (LA: top, BLA: middle and PIR: bottom) using a SVM classifier on the second day of social reward training.

**K)** The fraction of units with significant responses during the sensory period changes in response to learning on day2 (change in S+ responsive: LA = $\chi2(1, N=178) = 6.01$, *$p=0.014$; BLA = $\chi2(1, N=214) = 6.21$, *$p=0.013$; PIR = $\chi2(1, N=490) = 1.92$, $p=0.17$, change in S- responsive: LA = $\chi2(1, N=178) = 0.05$, $p=0.82$; BLA = $\chi2(1, N=214) = 0.92$, $p=0.34$; PIR = $\chi2(1, N=490) = 0.0$, $p=1.0$)

**L)** Correlation of the pre- and post-learning population activity during the sensory sampling period on day2.

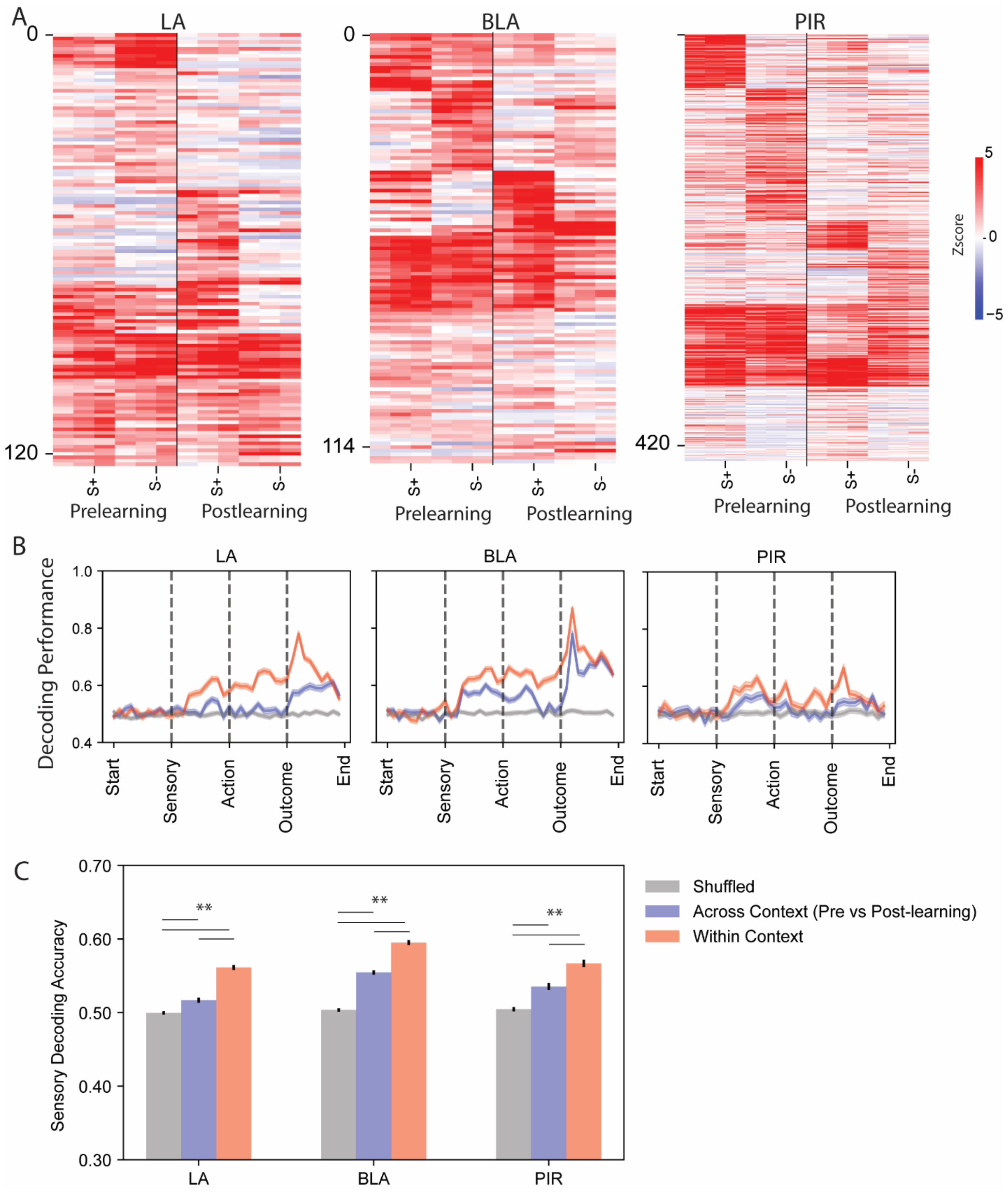


**Fig S4. Comparing neuronal responses before and after social reward learning.**

**A)** Z-scored single-neuron firing activity for the S+ and S- before and after social reward learning. The neuronal activity from each area was sorted using hierarchical clustering and only the clusters showing increasing firing to the stimuli are shown.

**B)** Using SVM classification, we tested whether population neuronal activity across contexts (pre- vs. post-learning) could accurately predict conspecific identity (S+ vs. S-). For the across context condition, we trained an SVM classifier on data from either pre or postlearning conditions and tested on the opposite dataset. Within context is the average of the SVM classifier performance trained and tested on data from before or after learning. We ran 100 iterations of each condition (shuffled, across context, within context) from the same 50 randomly selected neurons.

**C)** We compared the decoding performance of the three groups during the sensory epoch (shuffled labels, across context and within context) using a repeated-measures 1-way ANOVA and follow-up t-tests. (**$p<0.01$).

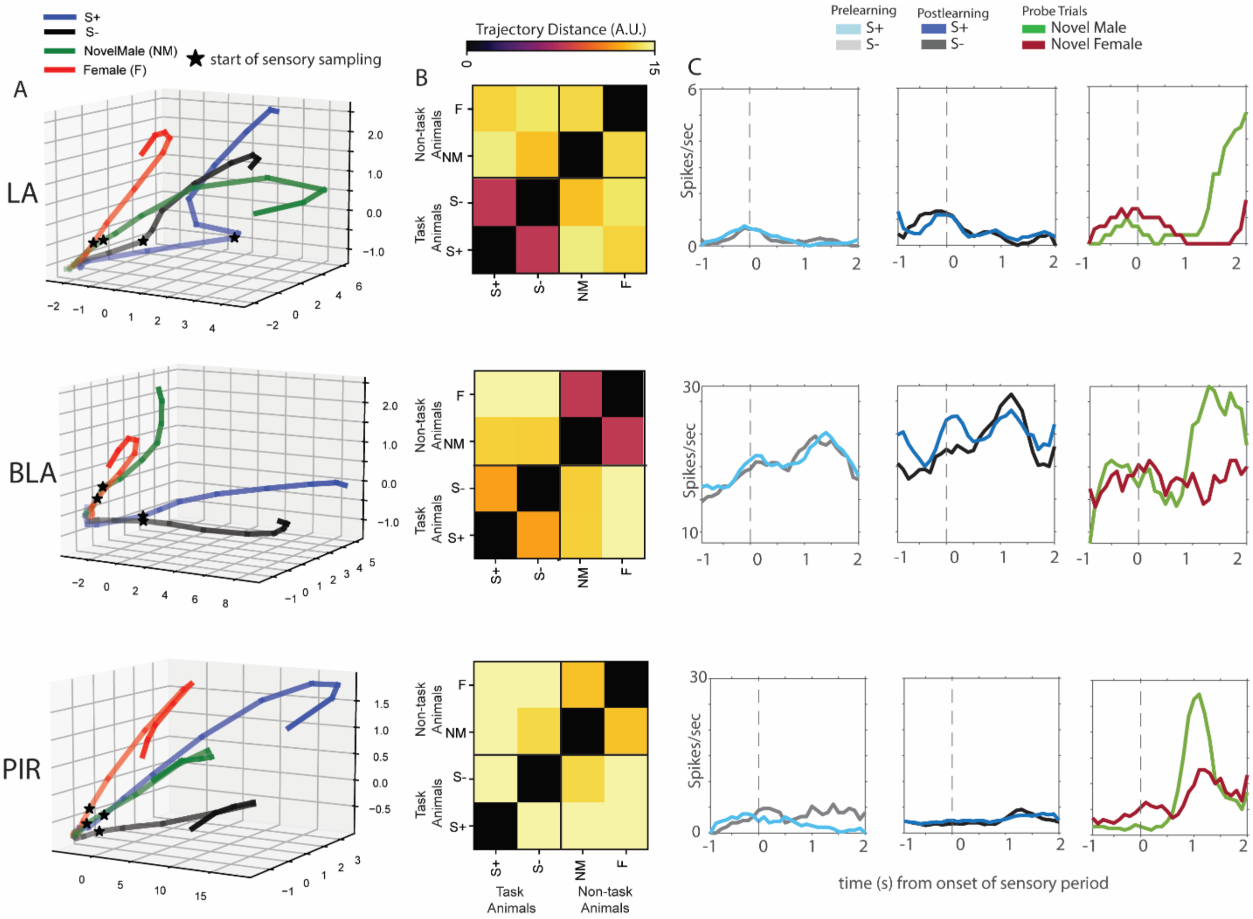


**Figure S5. Representation of probe trial animals in the LA, BLA and PIR.**

(**A**) Neuronal trajectories calculated from population activity in the LA, BLA and PIR reveal distinct representations of the task and non-task animals during the sensory period (sensory period onset indicated with a black star).

**B)** Quantification of the trajectory distance illustrates the diverging representations of task (S+ and S-) and non-task (NM and F) animals in the LA and BLA (top and middle). In contrast the PIR (bottom) has distinct representations of all 4 animals.

**(C)** Representative single-units from the LA, BLA and PIR (top, middle and bottom) that show strong responses to the novel male.

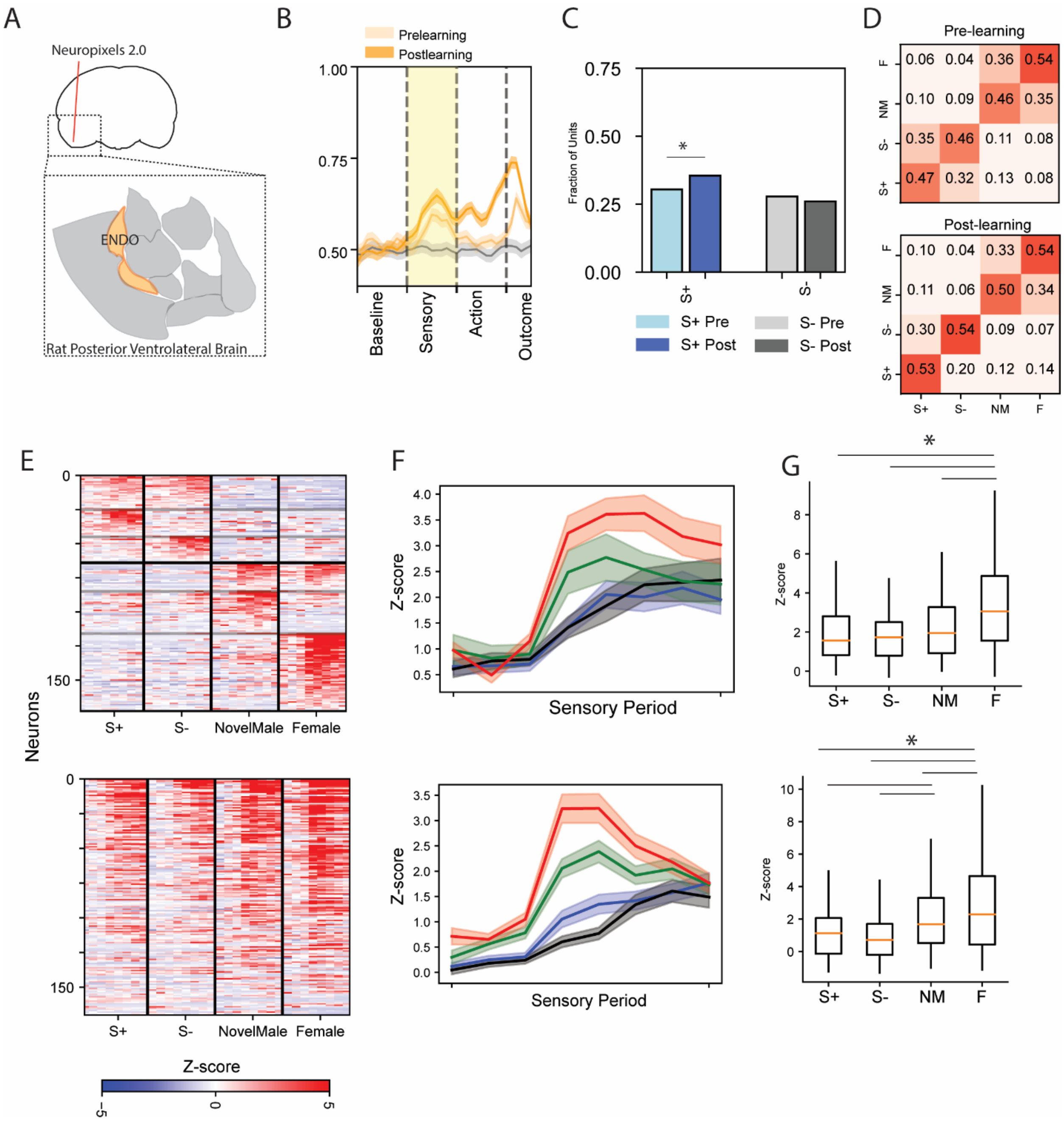


**Fig S6 Social identity, reward and familiarity in the endopiriform cortex.**

**A)** Alongside recording from the LA, BLA, and PIR, we recorded from large populations of neurons in the endopiriform cortex (ENDO, N=795).

**B)** Stimulus identity could be reliably decoded from ENDO population activity and performance increases slightly with social reward learning.

**C)** ENDO neurons show changes in response to social reward learning. The fraction of units responding to the S+ increases significantly with learning (change in S+ responsive: ENDO = $\chi2(1, N=795) = 4.33$, $*p<0.05$; change in S- responsive: ENDO $\chi2(1, N=795) = 0.54$, $p=0.46$)

**D)** Decoding of the identity of the 4 conspecifics before and after learning.

**E)** Z-scored firing activity for single neurons during the sensory period. The ENDO contains single-neurons that are selective for each of the four animals (top) and neurons that show general increases in activity to all conspecifics (bottom).

**F)** The selective neurons (top) and general neurons (bottom) have stronger firing activity for the non-task animals, with a specific increase in the response to the female conspecific.

**G)** Quantification of the differences in z-scored firing activity during the peak sensory period for selectively tuned ENDO (top, *$p<0.05$, 1-way ANOVA with post-hoc t-tests) and neurons with general selectivity (bottom, *$p<0.05$, 1-way ANOVA with post-hoc t-tests).

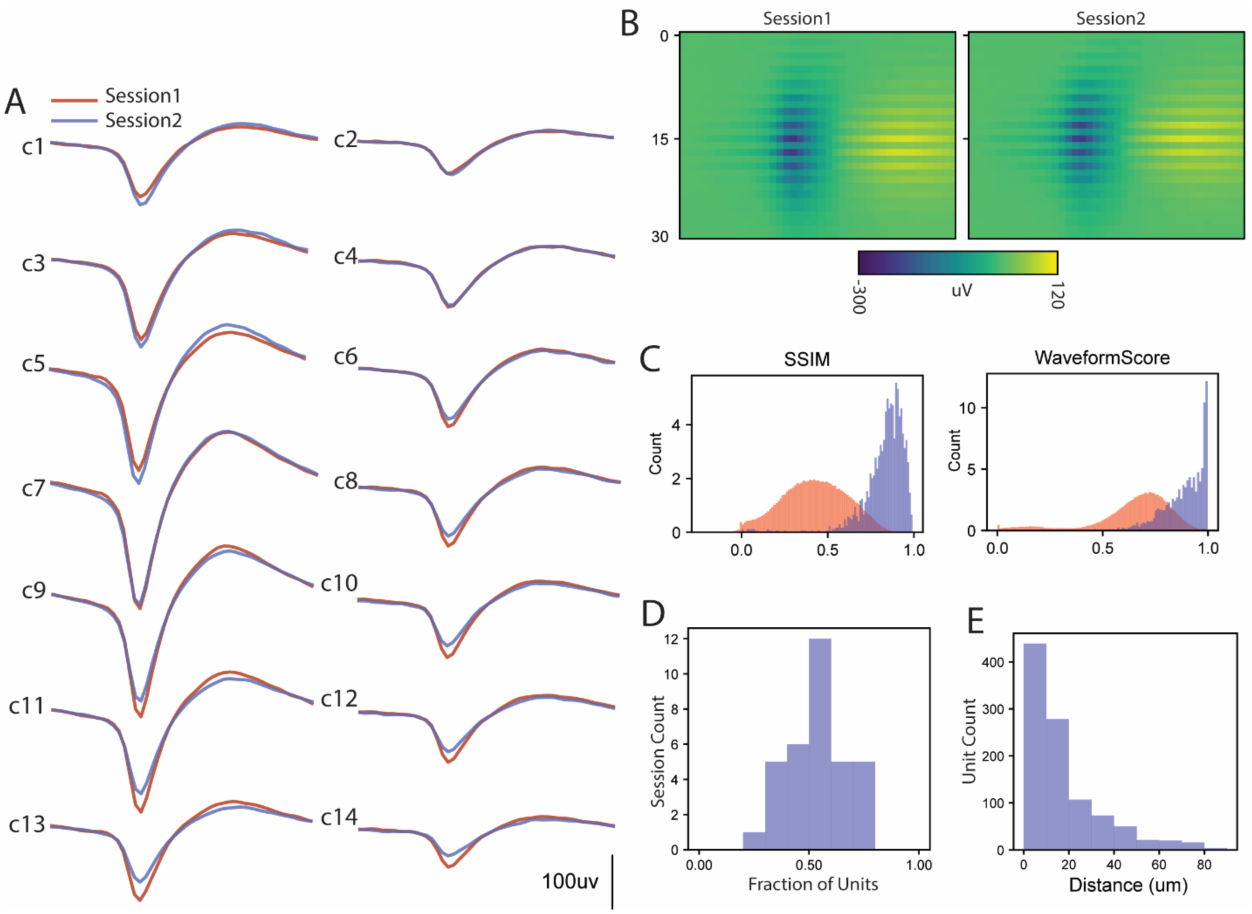


**Fig S7 Unit-tracking between same-day task recording and naturalistic social behavior.**

**A)** A representative single-unit that was identified both during session1 (social reward task) and session2 (naturalistic social behavior). On average there was a 1-2h gap between the two sessions.

**B)** Leveraging the high-density placement of electrodes on Neuropixels probes, we calculated similarity metrics from the waveform heatmap to identify the same units across sessions.

**C)** Two example metrics used for waveform matching including the structural similarity index measurements (SSIM) from the normalized waveform heatmap pictured in (B) and the waveformscore from Unitmatch calculations[1]. Successfully tracked units are in blue and adjacent neurons (<50um distance) are in red.

**D)** The fraction of units that were successfully tracked between the two same-day sessions (social reward task and naturalistic social behavior, 0.54 ± 0.14 mean ± standard deviation).

**E)** Unit depth was stable across the two sessions. We calculated the distance or drift of each individually tracked unit across. (14.61 ± 23.34um, mean ± standard deviation, electrode row distance = 15um).

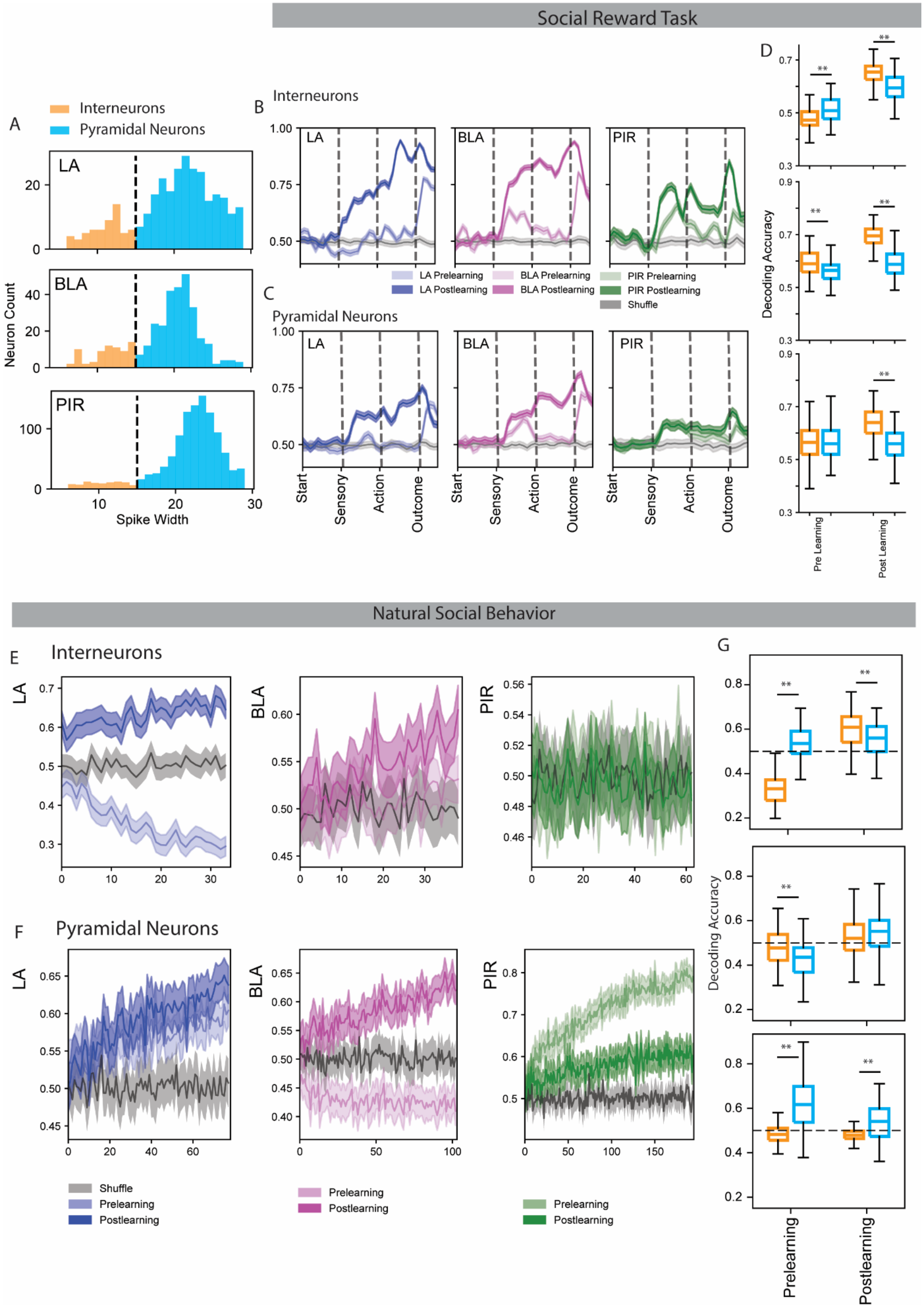
Social Reward Task
A
Interneurons
Pyramidal Neurons
LA
BLA
PIR
Neuron Count
Spike Width
B
Interneurons
C
Pyramidal Neurons
LA Prelearning
LA Postlearning
BLA Prelearning
BLA Postlearning
PIR Prelearning
PIR Postlearning
Shuffle
Start
Sensory
Action
Outcome
D
Decoding Accuracy
Pre Learning
Post Learning
Natural Social Behavior
E
Interneurons
F
Pyramidal Neurons
Shuffle
Prelearning
Postlearning
G
Decoding Accuracy
Prelearning
Postlearning

**Fig S8 Differential effects of pyramidal neurons and interneurons across the 3 regions.**

**A)** We used the spike width of the neuronal waveform to divide neurons into two groups (putative pyramidal and putative interneurons) and then calculated how each class of neurons contributes to social coding.

**B-C)** Across the 3 regions, interneurons have an outsized effect of the learning of social reward compared to pyramidal neurons. We still see pre-learning identity encoding in the BLA and PIR with both interneurons and pyramidal neurons.

**D)** Quantification of decoding performance during the sensory period from 50 neurons (100 iterations). Two-way ANOVAs revealed significant main effects of neuron type, learning condition and their interaction for the LA ($F$s > 6.8, $p$s < $9.5\times10^{-3}$), BLA ($F$s > 59.8, $p$s < $1.7\times10^{-13}$), and PIR ($F$s > 30.3, $p$s < $7.9\times10^{-8}$). Post-hoc comparisons showed significant differences between interneuron and pyramidal neuron decoding accuracies in the LA, BLA, and in post-learning PIR data. N=100 iterations.

**E-F)** Across the task and naturalistic social behavior contexts, we calculated the contribution of the two neuronal classes. We trained an SVM classifier on either interneuron (E) or pyramidal neuron (F) sensory task data with increasing numbers of neurons and tested on the first 20s of naturalistic social behavior

**G)** Two-way ANOVA results show significant main effects of neuron type, learning phase, and their interaction for LA ($F$s > 96.4, $p < 7\times10^{-20}$), BLA ($F$s > 7.5, $p < 6.6\times10^{-3}$), and PIR ($F$s > 25.3, $p < 8.4\times10^{-7}$). Post hoc comparisons revealed significant pairwise differences for specific neuron type and learning phase combinations (**p<0.01).

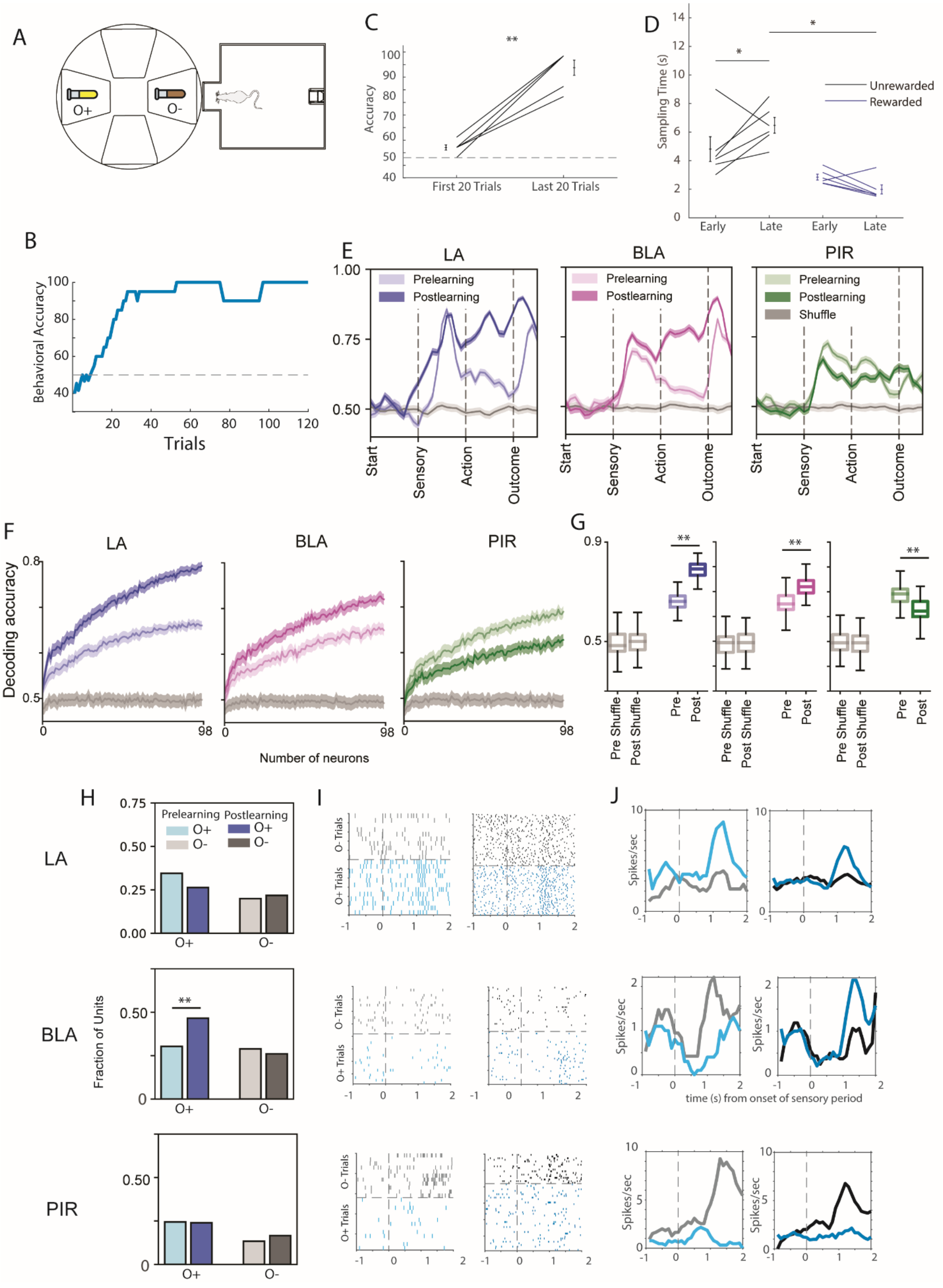
A
O+
O-
C
**
Accuracy
First 20 Trials
Last 20 Trials
D
Sampling Time (s)
Unrewarded
Rewarded
Early
Late
B
Behavioral Accuracy
Trials
E
LA
BLA
PIR
Prelearning
Postlearning
Shuffle
Start
Sensory
Action
Outcome
F
Decoding accuracy
Number of neurons
G
Pre Shuffle
Post Shuffle
Pre
Post
H
Fraction of Units
I
O- Trials
O+ Trials
J
Spikes/sec
time (s) from onset of sensory period

**Figure S9. Non-social odors are strongly represented in the amygdala and piriform.**

**A)** Implanted rats were trained on a variant of task where the S+/S- conspecifics were replaced with odor extracts (e.g. vanilla and almond, O+/O-).

**B)** Rats were able to learn the assigned valence of the odors within a single day's session. (46±9 trials before learning, mean±SEM, N=6 sessions).

**(C)** Accuracy increased in the olfactory task from the first 20 to the last 20 trials (N=6rats, accuracy first 20 trials: 55±1.29, accuracy last 20 trials: 94.16±3.74, paired t-test, **p<0.01)

**D)** Average sampling times are shown for the first 20 and last 20 trials. A mixed RM-ANOVA was conducted with trial (early vs. late) as a within-subjects factor and reward status (rewarded vs. unrewarded) as a between-subjects factor to examine their effects on sampling time. The main effect of trial was not significant (F(1,10)=0.598, p=0.457, but the interaction effect (trial x reward status) was significant F(1,10)=6.014, *p<0.05. Post-hoc comparisons (Bonferroni-corrected indicated that sampling time was higher in unrewarded compared to rewarded presentations in later trials.

**E)** The O+/O- could be reliably decoded from both the pre- and post- learning data across the 3 regions. Decoding performance was high in the pre-learning data suggested robust odor identity encoding in all three regions.

**F-G)** We quantified the odor decoding during the sensory period by training a linear SVM classifier with increasing numbers of neurons per region and quantifying the decoding performance with the maximum number of neurons (98). All regions show substantial ability to decode odor identity compared to shuffled data. A repeated-measures ANOVA revealed significant main effects of learning conditions F(3,2486)=9002, **p<0.01) and brain region, F(2, 2486)=372 **p<0.001 as well as a significant condition x region interaction F(6,2486)=477. Post-hoc comparisons showed significant learning increases in the LA (*pre:* 0.66±0.003 *post:*0.79±0.003*)* and BLA *(pre:*0.65±0.005 *post:*0.72±0.004*)* and a significant decrease with learning in the PIR (*pre:*0.69±0.004 *post:*0.63±0.004*)*.

**H)** From the onset of the task, the LA more strongly represented the O+ and the representation of the O- increased with learning, while the BLA showed increased representation of the O+ with learning. The piriform does not show any consistent changes with odor learning. (change in O+ responsive: LA = $\chi 2$(1, N=110) = 1.37, p=0.24; BLA = $\chi 2$(1, N=142) = 5.43, *p<0.05; PIR = $\chi 2$(1, N=217) = 0.0, p=1.0, change in O-responsive: LA = $\chi 2$(1, N=110) = 0.03, p=0.86; BLA = $\chi 2$(1, N=142) = 0.16, p=0.69; PIR = $\chi 2$(1, N=217) = 0.65, p=0.42)

**I-J)** Example representative units for the 3 areas (LA: top, BLA: middle, PIR: bottom).
